# Diversity of motility gene signatures distinguishes commensal Clostridia that elicit divergent host immune responses

**DOI:** 10.1101/2025.07.18.665615

**Authors:** Lennard W. Duck, Melissa S. Jennings, Jung-Shan Hsu, Covenant F. Adeboboye, E. Leighann Morgan, Kiarra J. Coger, Barbara J. Klocke, Dave D. Hill, Katie L. Alexander, Alexander F. Rosenberg, Goo Lee, Qing Zhao, Charles O. Elson, Craig L. Maynard

## Abstract

Adaptive immune responses to commensal flagellins are hallmarks of Crohn’s disease, but it is unclear whether flagellins themselves promote inflammation or whether flagellated commensals can also be colitogenic. Here, we show that the arrangement of motility loci and the diversity of encoded flagellins can separate flagellated gut-derived Clostridia into at least 2 functionally distinct groups. In gnotobiotic mice, both groups induce tolerogenic responses but only one group promoted tissue inflammation following barrier disruption. Accordingly, specific flagellins expressed by members of this pro-inflammatory group displayed a heightened capacity for TLR5 activation which could be altered by modification of a defined region of the flagellin D0 domain. Finally, bacteria belonging to the pro-inflammatory group were found to be elevated in Crohn’s disease biopsies. Collectively, our study identified key features of specific commensal bacteria that possess colitogenic potential and revealed one mechanism whereby these organisms can potentially initiate intestinal inflammation.

## INTRODUCTION

Taxonomically related gut bacteria, including different strains of the same species, can have divergent effects on the host. In many cases, these differences are due to differential expression of specific genes and microbial pathways. What is less appreciated is the importance of differences within similar gene products in predicting or perhaps dictating the outcomes of host-microbe interactions, which may underpin the etiology and/or progression of gut inflammation. Within the broader gut metagenome, cell motility is a minimally represented function (*1*) mediated by propulsive structures such as flagella and fimbriae (*2*). However, in specific niches of the colon, including among Clostridia species that are dominant producers of short chain fatty acids (SCFAs), flagella-dependent motility is a highly represented function. Accordingly, humans and mice express cell surface (*3*), cytosolic (*4*), and, secreted (*5*) proteins that recognize flagellin – the basic monomeric unit of flagella. The genomes of most flagellated Clostridia harbor multiple flagellin genes but the extent to which each is expressed, and the impact of this diversity on host-commensal interactions in health and disease, are not clear.

Flagella-dependent motility is a feature of several species of Clostridia that colonize the intestine. These bacteria encode varying numbers of flagellins – the basic subunits of flagella - and the machinery necessary for flagella synthesis and function. The flagellin monomer is comprised of generally well-conserved α-helical N- and C-terminal regions that polymerize within the inner core of the flagellar filament to form D0 and D1 domains. The intervening hypervariable regions of each flagellin are arranged into turn-folded β-sheets that form the external facing and often heavily glycosylated regions of flagella. Flagellar-dependent motility requires the synthesis and assembly of flagellin monomers to form flagellar filaments that are then powered by a complex system of motor proteins. The genes responsible for these processes are often found in confined stretches of the bacterial genome in generally well-conserved motility loci. Interestingly, a previous study identified and characterized 3 motility loci - *flgB*-*fliA*, *flgM*-*flgN*/*fliC*, and *mbl*-*flgJ* - in the genomes of select *Lachnospiraceae* (*6*). The *mbl*-*flgJ* domain, comprised of *mbl*, *flgF*, *flgG*, and *flgJ*, existed in *Lachnospiraceae* but not the *Ruminococcaceae* genome studied and is separated by hundreds of genes from the larger, generally well-conserved *flgB-fliA* locus. Importantly, in the *Ruminococcaceae* genome, the genes *flgF* and *flgG*, important in assembly of the flagellar rod, were present within the *flgB-fliA* locus, meaning that those functions are maintained despite the distinct gene order. This divergence was attributed to the taxonomic distance between *Lachnospiraceae* and *Ruminococcaceae* but also raises the question as to whether differences in motility gene order can be predictive of functional and/or behavioral differences even among taxonomically similar bacteria.

Flagellins or flagellin-like structures are encoded by most major phyla of archaea and bacteria. Among human commensal bacteria, flagellins are predominantly encoded by bacteria that belong to the phyla Pseudomonadota (Proteobacteria) and Bacillota (Firmicutes) (*7*). Much of our knowledge of flagellin function in the gut has been obtained through the study of Proteobacteria, particularly *Escherichia coli* and pathogenic *Salmonella* sp. However, among human gut commensals, bacteria of the *Lachnospiraceae* family within the order Lachnospirales and the phylum Bacillota (Firmicutes) are the dominant, albeit not exclusive, expressors of flagellins (*8*). Some *Lachnospiraceae* flagellins are believed to well be tolerated by the host in part via a mechanism described as “silent recognition”, predicated on hypoactivation of the cell surface flagellin receptor, toll-like receptor 5 (TLR5) despite normal binding (*8*). Additionally, flagellins from *Lachnospiraceae* bacteria *Roseburia hominis* (*R. hominis*) and *R. intestinalis* can promote induction and/or expansion of regulatory T cells (Treg cells) (*9, 10*), likely facilitating the co-existence of these organisms with the host. These tolerogenic mechanisms stand in stark contrast to the robust TLR5 activation mediated by pathogen-derived flagellins including the dimerization dependent hyperactivation of TLR5 by flagellin from *Salmonella* (*8*). On average, *Lachnospiraceae* are reduced in abundance in Crohn’s disease (*11–13*), supporting the notion that loss of this regulatory axis is causally associated with gut inflammation. On the other hand, elevated serum anti-flagellin IgG and elevated flagellin-reactive effector T cells are common in Crohn’s disease and reactivity to multiple *Lachnospiraceae* flagellins is associated with worse patient outcomes (*14*). These divergent observations suggest that immune responses to distinct flagellins or flagellated bacteria can potentially tip the balance between homeostasis and immune activation. This concept seems even more plausible given the diversity of flagellated organisms and the intimate association of *Lachnospiraceae* and other Clostridia with mammalian hosts (*15*).

Based on the foregoing information, we reasoned that motility gene signatures may represent a suitable starting point from which to begin to delineate potential differences among motile commensal bacteria that may underpin differential colitogenic capabilities and may even transcend taxonomy. Here, we show that the relative position of specific motility loci overlaps with diversity of encoded flagellins to distinguish certain flagellated gut commensals which we collectively refer to as Group1 (G1) and G2 bacteria. Whereas G1 bacteria encode a diverse set of flagellins including one resembling previously described “silent” flagellins, G2 bacteria encode a more homogenous set of flagellins. Colonization of mice with representative G1 or G2 isolates resulted in the accumulation of Foxp3+ RORγt+ Treg cells in the colonic lamina propria, and colonization-dependent IgA responses, both of which facilitate host-microbiota mutualism. However, following mucus layer disruption, G2-, but importantly not G1-colonized mice developed colonic inflammation. Moreover, representative G2-encoded flagellins activated TLR5 and triggered downstream cascades more robustly than G1-encoded flagellins. Furthermore, the stimulatory capacities of representative G1- or G2-encoded flagellins could be significantly altered by modifying a defined region of the D0 domain. Interestingly, analysis of publicly available microbiota sequencing data revealed that certain G2 but not G1 bacteria are increased in ileal biopsies of CD patients. Collectively, our data demonstrate that flagellated commensals resident in the intestine include functional consortia with distinct capabilities for immune activation, and identify potential candidates for future exploration of commensal-driven inflammation.

## RESULTS

### Separation of Clostridia genomes based on motility gene positioning and flagellin diversity

We started with 108,290 genomes in the class Clostridia identified in the National Center for Biotechnology Information (NCBI) database. Using the Pathosystems Resource Integration Center (PATRIC) database this list was computationally narrowed to 759 representative genomes (**Fig. S1**). We then manually selected 101 genomes that are gut-derived and encode at least one flagellin. We and others previously isolated flagellated Clostridia from the mouse cecum (*16*), human feces (*17*), and human intestinal biopsies (NCBI BioProject 28331) (*18*) and inclusion of these genomes brought the total number to 163 genomes of flagellated Clostridia, including 96 human and 20 murine isolates, spanning 4 Orders – Christensenellales, Eubacteriales, Lachnospirales, and Peptostreptocaccales – and 12 families, and harboring a total of 518 flagellin genes (**Fig. S1, Table S1**). We first aligned the motility loci previously identified by Neville et al. (*6*) to determine whether there was any divergence in loci positioning, similar to what was noted for *Lachnospiraceae* versus *Ruminococcaceae*. As predicted, 59 genomes, the majority of which were *Lachnospiraceae*, contained the 4-gene locus *mbl*, *flgF*, *flgG*, and *flgJ* but the remaining genomes did not, although a fraction had the *flgF* and *flgG* genes present in the larger *flgB*-*fliA* locus (**Fig. 1A**). Within the *flgB*-*fliA* locus, the *flhA* gene was present and highly conserved across all genomes and thus we used the number of genes separating *flhA* from *flgF* as a measure of the proximity of these 2 loci (**Fig. 1A**). In all *mbl*-*flgJ*-containing genomes, this *flhA to flgF* distance ranged from 120 - 5854 genes with a mean distance of 1310 genes (**Fig. 1B**). Thirty-six genomes comprised of mostly Clostridiaceae displayed *flhA to flgF* distances ranging from 3 – 8 genes with a mean distance of 6.6 genes. In 65 of the remaining 68 genomes, the *flhA* and *flgF* genes were separated by a single gene. Genomes 162 and 163, despite passing our selection based on their gut origin and encoding of flagellin genes, appear to have lost their motility genes, but were kept and grouped with genomes 96-161 based on having a motility gene distance of 1 or lower. Altogether, this group was highly diverse but dominated by *Lachnospiraceae* and *Oscillospiraceae* (**Fig. 1B**). Thus, there is distinct spatial arrangement of flagella-related motility genes among flagellated Clostridia from human and non-human hosts independent of taxonomy at the family level.

**Fig. 1.**
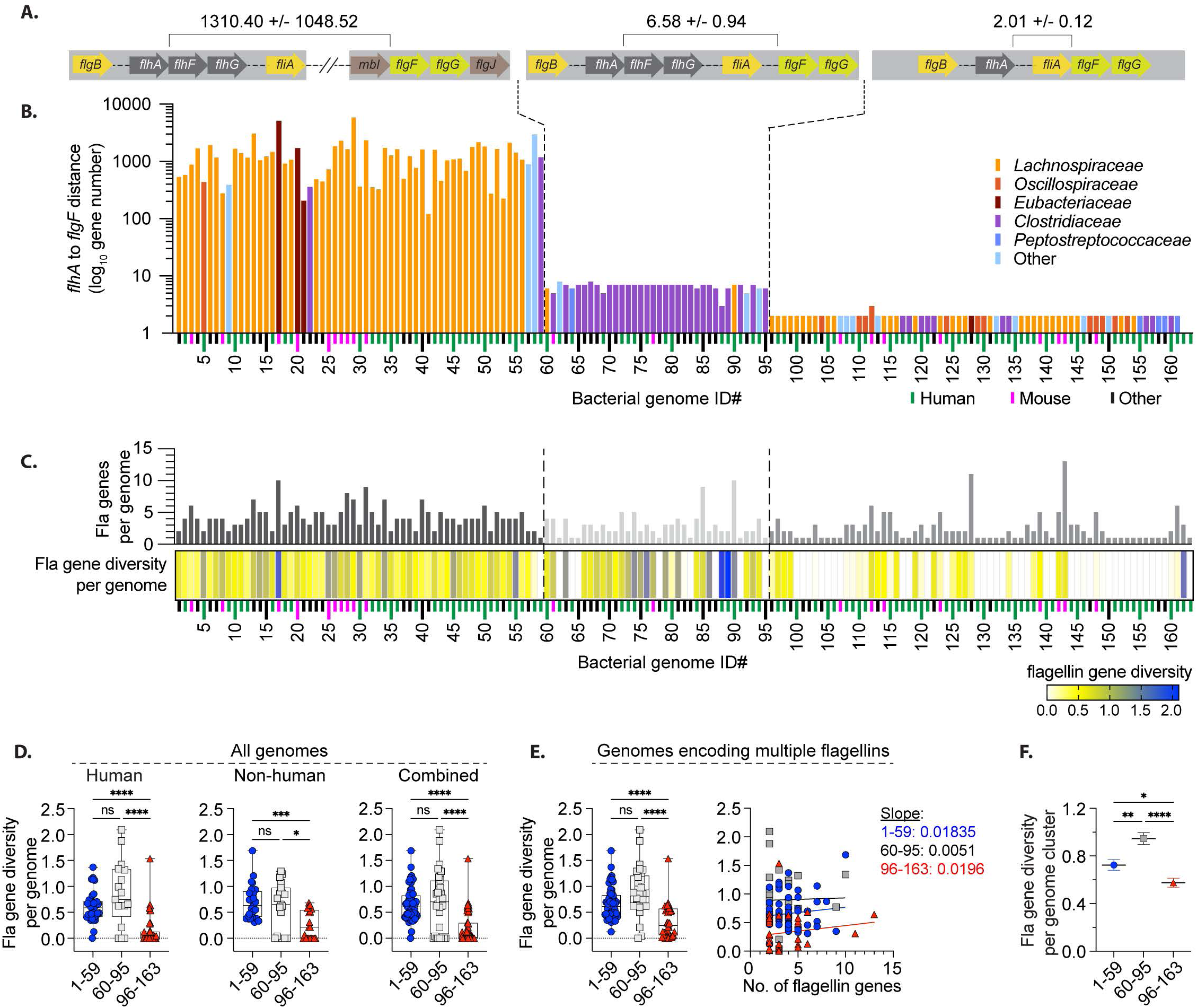
Separation of representative Clostridia genomes into distinct groups based on motility loci positioning and flagellin diversity. (A). Schematic representation of the motility loci examined in the 163 Clostridia genomes (x-axis, detailed in Table S1) studied to determine the *flhA* to *flgF* distance. (B) Graph shows the calculated *flhA* to *flgF* distance of the 163 genomes ordered based on the 3 distance categories and color-coded by family. Distances represent the difference between PATRIC PEG numbers for *flhA* and *flgF* in all the genomes studied. Colored markers on the x-axis indicate isolates derived from human (green), mice (pink), or other species (black) shown in Table S1. (C) Bar graph (upper) shows total number of flagellin genes encoded by each genome while the heatmap (lower) indicates the mean number of amino acid substitutions at each position for all flagellins encoded in each genome. (D) Graphs show flagellin diversity score for each genome derived from human and non-human sources or for the total collection of 163 genomes and grouped based on the *flhA*-*flgF* distances shown in 1B. (E) After genomes encoding a single flagellin were excluded, flagellin diversity values for remaining genomes were plotted against genome cluster (left) or against total number of flagellin genes (right). Slope values are shown that support the lack of correlation between flagellin gene number and diversity within the genome clusters. (F) Graph shows the mean diversity of flagellins within each genome cluster and bootstrap (1000) generated SEM for each cluster. Graphs in D-F show mean +/− SEM. *p<0.0332, **p<0.0021, ***p<0.0002, ****p<0.0001

We then asked whether there is any relationship between the observed motility gene arrangement and the number and diversity of encoded flagellins. We found that among genomes 1-59 that feature multiple motility loci, 58 of 59 (98%) encode multiple flagellins. In contrast, among genomes 96-163, 38 of 68 (56%) encode multiple flagellins (**Fig. 1C**, upper). We also assessed flagellin diversity within each of the 163 genomes by calculating the number of substitutions per site across 208 amino acid positions in each of the flagellins encoded by that genome (**Fig. 1C**, lower). When considered as a group, genomes 1-59 displayed an average flagellin gene diversity that was significantly higher than that of genomes 96-163, whether the isolates were derived from human or non-human sources (**Fig. 1D**). Even after we eliminated the genomes that encode a single flagellin and therefore have a diversity score of 0, there was still a significant difference in flagellin diversity between genomes 1-59 and 96-163 (**Fig. 1E**, left). In fact, across all genome clusters, average flagellin diversity was more closely correlated with the genome cluster than with the number of flagellins encoded by each individual genome within that cluster, with line slopes approaching 0 (**Fig. 1E**, right). We also assessed the diversity of all flagellins within each genome cluster independent of the genome from which they were derived (**Fig 1F**) and observed significant differences between all 3 genome clusters. Importantly, cluster that includes genomes 60-95 displayed the highest within-group diversity, suggesting that individual flagellins from a given organism were less likely to be representative of the other flagellins in a given genome or in the entire cluster. This, coupled with the fact that this cluster appears to include subsets that bear resemblance to either of the other 2 genome clusters (Figs. B-E), we hesitated to consider this a coherent group resulting in exclusion of these genomes from further group-based analyses in this study. Collectively, these data show that the arrangement of motility loci and the diversity of the encoded flagellins combine to identify 2 highly distinct clusters of flagellated Clostridia independent of host species or any taxonomic relatedness particularly at the family level. For simplicity, we have collectively referred to genomes 1-59 as Group1 (G1) and genomes 96-163 as G2. G1 is primarily comprised of members of the *Lachnospiraceae* family whereas G2 has more diverse representation from multiple families with *Lachnospiraceae* and *Oscillospiraceae* being the most highly represented (**Fig. S2**).

### Both G1 and G2 bacteria colonize the colon mucus layer and induce tolerogenic responses at steady state

We previously isolated multiple species of commensal bacteria from the cecum of mice and demonstrated that they expressed flagellins that were recognized by antibodies in Crohn’s patient sera (*16*). Some of these isolates were included in the 163 genomes in figure 1 and somewhat surprisingly, even these isolates separated into either G1 or G2 based on our classification. We combined this biorepository of well-characterized anaerobic commensals that are readily culturable in our hands, with our well-established mouse models, to conduct *in vivo* studies aimed at determining the relative functional potential of G1 and G2 bacteria. We utilized our IL-10 transgenic reporter (10BiT) model (*19*) to generate gnotobiotic cohorts each harboring a trio of select G1 or G2 bacteria. For G1, we selected the *Roseburia*-like *Eubacterium* sp. 14-2, *Lachnospiraceae* bacteria 28-4 and A4, which correspond to genomes #20, #26, and #29, respectively. Considering the family-level diversity seen in G2, we utilized *Anaerotruncus* sp. G3, *Lachnospiraceae* bacterium M18-1 and *Oscillobacter* sp. 1-3, which correspond to genomes #112, #142, and #148, respectively. Germ-free females were colonized with respective G1 and G2 bacteria then bred to generate the progeny that would become the experimental animals and also used for propagation of the gnotobiotic cohorts (**Fig. S3A**). Therefore, all experimental animals acquired the bacteria via vertical transmission from the mothers, which was confirmed by polymerase chain reaction (PCR) on fecal bacterial DNA (**Fig. S3, B-C**). All G1 and G2 bacteria were detectable in mucosal scrapings from the cecum, as well as the proximal, middle, and distal colon. Importantly, both G1 and G2 bacteria were approximately 10-times more abundant in the proximal relative to the distal colon (**Fig 2, A-B**), in agreement with previous reports of enrichment of bacteria including *Lachnospiraceae* and *Ruminococcaceae* in the interfold regions of the proximal colon (*15*).

**Fig. 2.**
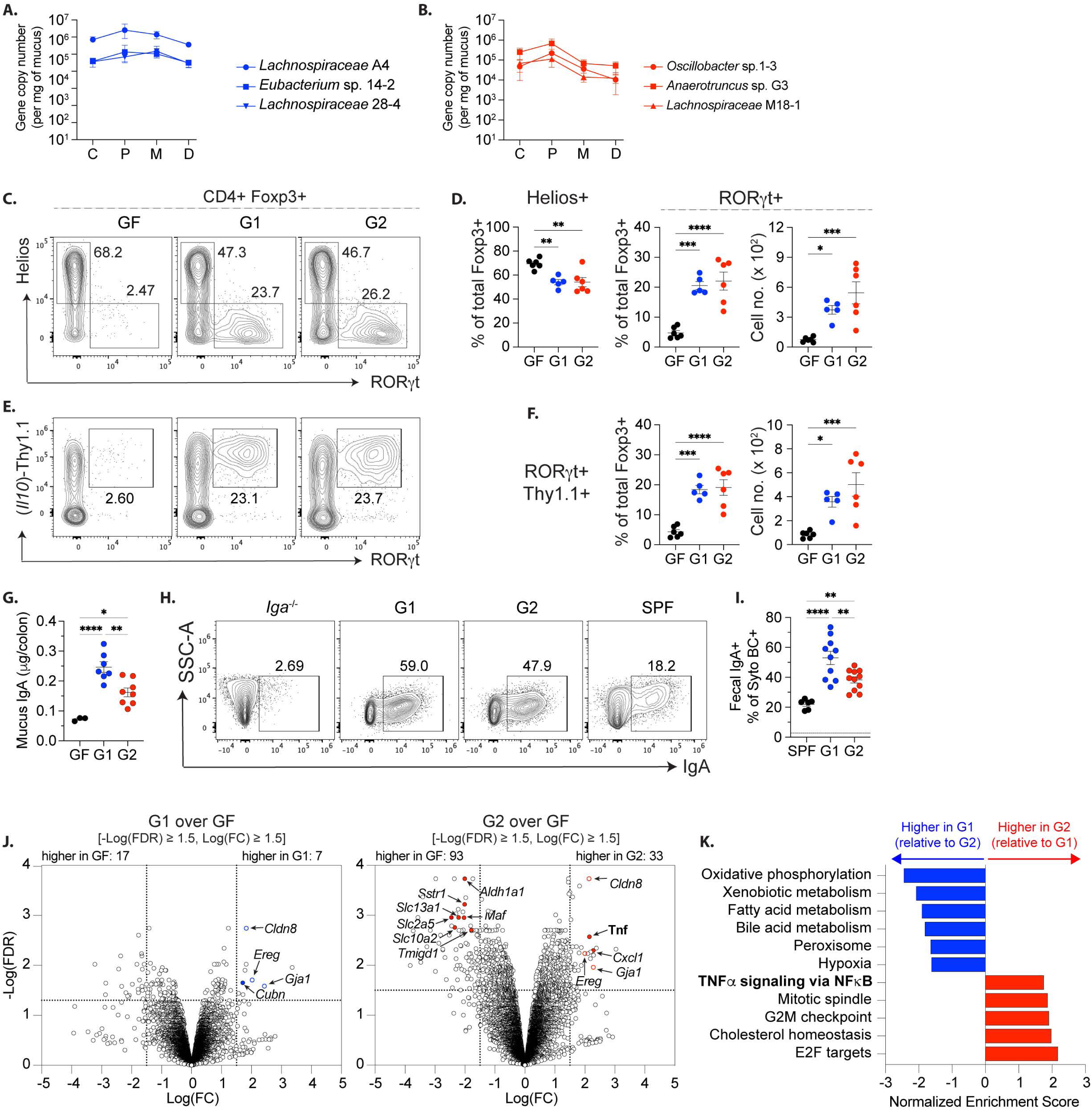
Host response to G1 and G2 bacteria at homeostasis. Graphs showing copy numbers of (**A**) G1 and (**B**) G2 bacteria detected by qPCR in the mucus layer of the cecum (C), proximal (P), middle (M) and distal (D) colon of gnotobiotic mice. (**C**) Gated CD4+ Foxp3+ cells were examined for expression of RORγt and Helios. (**D**) Graphs summarizing data collected as in (C) show frequencies and/or numbers of Helios+ and RORγt+ cells. (**E**) Gated CD4+ Foxp3+ cells were examined for expression of RORγt and Thy1.1. (**F**) Graphs summarizing data collected as in (E) show frequencies and/or numbers of RORγt+ Thy1.1+ cells. (**G**). Graph shows levels of IgA detected in colon mucus of 8-week-old GF, G1 and G2 mice. (**H**) FACS plots showing IgA coating of bacteria in the feces of G1, G2, and SPF mice with IgA-deficient mice included as negative controls. (**I**) Graph summarizing data collected as in H. Numbers in FACS plots indicate the percentage of cells in the respective gates. Results in A-I are pooled from at least 2 independent experiments. (**J**) Volcano plots illustrating average transcript expression by colonic epithelial cells from G1 or G2 mice (3 mice per group) compared to GF mice and determined by bulk RNA sequencing. Representative transcripts that were increased or decreased at least 1.5-fold in G1 or G2 relative to GF are indicated by colored open circles (similarly expressed) or colored closed circles (differentially expressed). (K) Graph shows transcriptional pathways more highly represented in colonic IEC of G1 mice relative to G2 mice (blue), and vice versa (red), under homeostatic conditions. Graphs in A, B, D, and F show mean +/− SEM. *p<0.0332, **p<0.0021, ***p<0.0002, ****p<0.0001.

Similar to the previously described ability of several Clostridia isolates (*20, 21*), colonization by both G1 and G2 bacteria resulted in significant accumulation of commensal-induced RORγt+ Helios-peripheral regulatory T cells (pTreg cells) in the colonic lamina propria (**Fig. 2, C-D**), the majority of which co-expressed immunosuppressive IL-10 (**Fig. 2, E-F**). In addition, colonized mice harbored elevated levels of colon mucus IgA that was generally more abundant in G1-relative to G2-colonized mice (**Fig. 2G**). Accordingly, we also detected significantly higher frequencies of IgA-coated fecal bacteria in G1 relative to G2 mice, both of which exceeded that of specific pathogen-free (SPF) mice (**Fig. 2, H-I**). Finally, we conducted bulk RNA sequencing of the colon epithelial cell fraction isolated from G1 and G2 mice and compared to the transcriptional signatures of age-and sex-matched germ-free mice. Overall, we detected approximately 5 times the number of transcripts that were at least 1.5-fold higher or lower in G2 bacteria versus G1 bacteria, suggesting that the G2 bacteria are generally more stimulatory.

In epithelial cells from both G1- and G2-colonized mice, we detected transcripts for genes including those encoding the integral junction protein claudin-8 (*Cldn8*), the gap junction protein connexin 43, also known as gap junction alpha-1 (*Gja1*), and the cell growth inducer epiregulin (*Ereg*). In contrast, transcripts of the chemoattractant C-X-C motif chemokine ligand-1 (*Cxcl1*), as well as tumor necrosis factor (*Tnf*) were significantly higher in G2 but not G1 mice (**Fig. 2J**). We also observed increased transcripts of the iron co-transporter cubilin (*Cubn*) in G1 mice while G2 mice showed reduced levels of transcripts for solute carrier family proteins (*Slc5a2*, *Slc10a2* and *Slc13a1*) as well as the transmembrane and immunoglobulin domain containing protein-1 (*Tmigd1*). Interestingly, reduced levels of *CUBN1*, *SLC5A2*, *SLC10A2*, *SLC13A1*, AND *TMIGD1* have all been associated with gut inflammation (*22–24*). In addition, comparative gene set enrichment analysis revealed higher expression of genes associated with mitochondrial function in G1 colonized mice, whereas G2 bacteria appear to activate pathways related to inflammation and cellular proliferation (**Fig. 2K**). Collectively, these data show that at homeostasis, both G1 and G2 bacteria establish mucosal niches aided by the induction of anti-commensal IgA and the accumulation of IL-10-producing pTreg cells. Particularly in G2-colonized mice, these regulatory mechanisms appear to counter the potential for epithelial hyperactivation by G2 bacteria, thereby helping to maintain immune homeostasis with this community.

### G2 bacteria induce colonic inflammation following disruption of the mucus layer

Based on the results in figure 2J, we next sought to qualitatively assess the impact of G1 and G2 bacteria on the colonic mucus barrier and to determine potential consequences of physical contact between both groups of bacteria and the colonic epithelium – mimicking an interaction that would likely occur in IBD. Gnotobiotic mice were left untreated or fed 1% dextran sulfate sodium (DSS) for 5 days (**Fig. 3A**) to disrupt the mucus barrier, potentially enabling the interaction of the bacteria with colonic epithelial cells. As controls, we utilized GF and SPF mice. Staining of the colon mucus layer with the mucus-binding lectin wheat germ agglutinin (WGA) indicated that G1 and G2 colonization resulted in a more robust mucus barrier comprised of recognizable inner and outer layers, in contrast to uncolonized GF mice (**Fig. 3B**, upper panel). In addition, G1 and G2 bacteria were found in close proximity to, and even intimately associated with the inner mucus layer of the respective gnotobiotic mice (**Fig. 3B**, upper panel). Importantly, following DSS exposure, both G1 and G2 bacteria were then present at the epithelial border in direct contact with epithelial cells (**Fig. 3B, lower panel**), confirming the DSS-mediated mucus erosion. Relative to G1 mice, G2 mice displayed elevated fecal levels of the anti-siderophore lipocalin-2 (Lcn2) (**Fig. 3C)**, an early indicator of pro-inflammatory activation in the gut that importantly, can be produced downstream of TLR5 stimulation (*25, 26*). Histological analysis of hematoxylin and eosin (H&E)-stained colonic tissue confirmed induction of moderate tissue inflammation and epithelial damage in DSS-treated G2, but not G1 mice (**Fig. 3, D-G**). Importantly, this response was moderate relative to the inflammation induced by a complete microbiota in SPF mice. We subjected the epithelial fraction of all 4 groups to semi-quantitative PCR analysis and found that unlike G1-colonized mice and relative to control-treated counterparts, epithelial cells from DSS-treated mice G2 mice showed statistically significant increases in transcripts for tumor necrosis factor (*Tnf*), interleukin-1 beta (*Il1b*), as well as the neutrophil chemoattractant CXC motif chemokine ligand 1 (*Cxcl1*) (**Fig. 3, H-J**). Collectively, these data demonstrate that G2, but not G1 bacteria can induce moderate colonic inflammation following mucus barrier disruption.

**Fig. 3.**
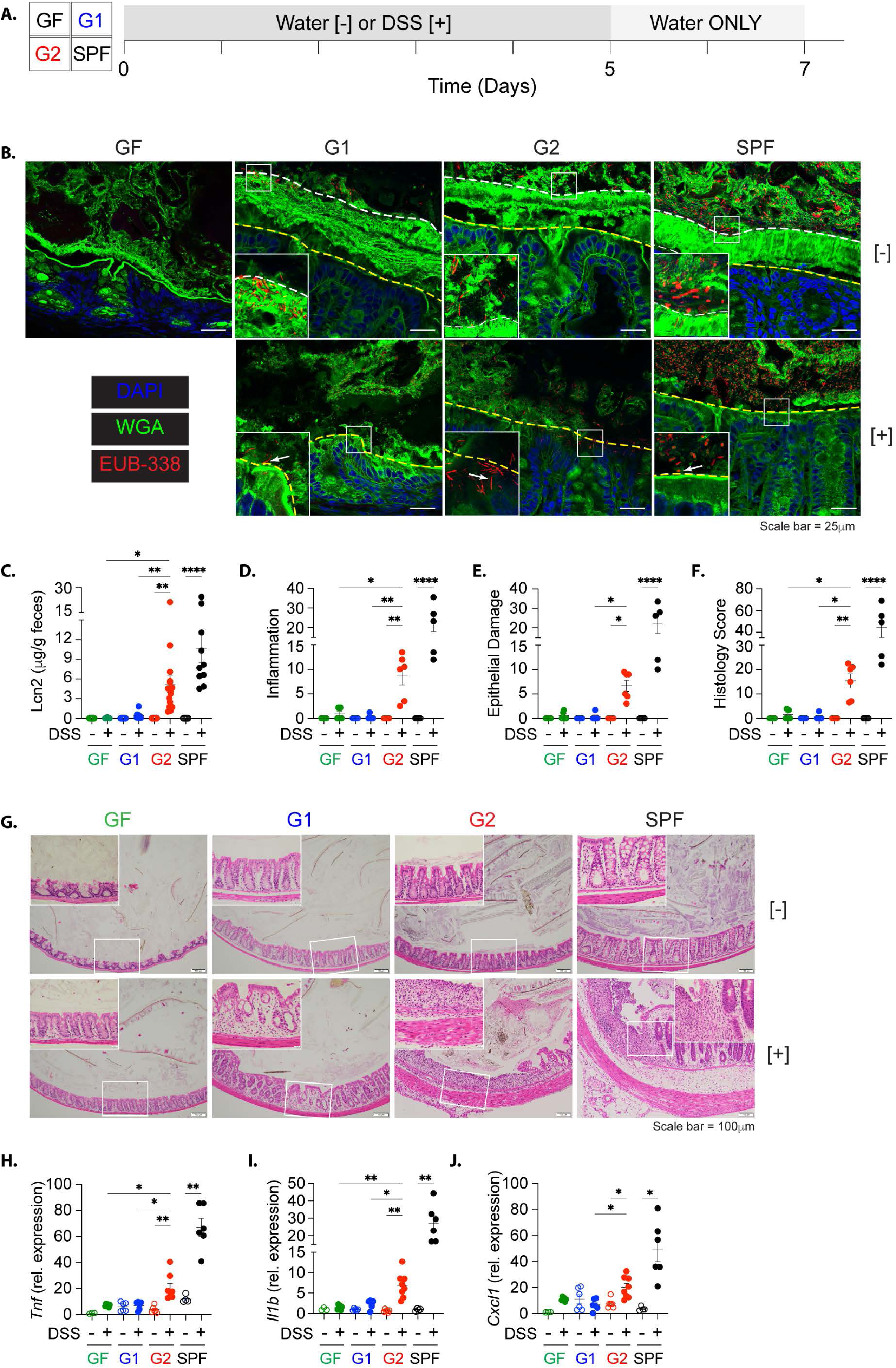
Host response to G1 and G2 bacteria following disruption of the colon mucus layer. (**A**) Overview of DSS experiments in gnotobiotic mice. (**B**) Immunofluorescence staining of Carnoy-fixed colon tissue from GF, G1, G2, and SPF mice fed normal drinking water (upper panel) or DSS (lower panel). Colon tissue was stained for epithelial cell nuclei (DAPI; blue) mucus (wheat germ agglutinin; green) and bacteria were detected by fluorescence in situ hybridization (EUB-338 universal bacteria probe; red). Dashed white line demarcates the border between inner and outer mucus layers while the dashed yellow line demarcates the epithelial border. Representative confocal images from one of 3-5 mice per group are shown at 60X magnification after examining at least 3 slides per mouse and at least 5 high-power fields per slide. (**C**) Fecal Lcn2 measured by ELISA and normalized to fecal weights. H&E-stained colonic tissue from control or DSS-treated mice were examined to assess (**D**) inflammation and (**E**) epithelial damage which were added to generate a histology score (**F**). (**G**) representative photomicrographs of H&E-stained tissue from all groups at 10X magnification, (inset = 20X). Total epithelial cells were enriched from the colon of the respective mice and qPCR was used to determine relative transcript levels of (**H**) *Tnf*, (**I**) *Il1b*, and (**J**) *Cxcl1*. All data are pooled from 2-3 experiments. Graphs show mean +/− SEM. *p<0.0332, **p<0.0021, ***p<0.0002, ****p<0.0001.

### Assessment of flagellin expression by representative G1 and G2 bacteria

Cxcl1 is known to be induced downstream of TLR5 (*27*). Therefore, the foregoing results suggested that flagellins may be at least partially responsible for the *in vivo* responses observed and that the differential ability to activate TLR5 may further distinguish G1 from G2 bacteria. To determine which flagellins to test in this setting, we generated pure cultures of the G1 and G2 bacteria used to colonize our gnotobiotic mice, and following mechanical disruption, performed liquid chromatography-mass spectrometry (LC-MS) on the enriched proteins. In all extracts of mouse G1 and G2 isolates, the most prevalent proteins were flagellins (**Fig. S4A**, **Tables S3-S8**). However, surprisingly, “FlaB-like” flagellins, which are only encoded by bacteria we classified as G1, were either not detected by our assay (*Lachnospiraceae* bacterium 28-4), or the signals were all at levels between 1% and 6% of the lowest expressed flagellins (*Lachnospiraceae* A4, *Eubacterium* sp. 14-2) comparable to the expression levels of intracellular proteins including the small ribosomal subunit protein uS5 (Rps2) (**Fig. S4, A-B and Tables S3-S5**). This was also the case when we analyzed extracts of human isolates *R. hominis*, *R. intestinalis* and *R. inulinivorans*, where *R*. *hom*inis FlaB, which was previously studied as the prototype “silent” flagellin (*8*), or other *Roseburia* FlaBs were either not detected (*R. intestinalis*) or detected at spectral values of 1-3% of the lowest expressed flagellins (*R. hominis*, *R. inulinivorans*) (**Fig. S4C, and Tables S9-S11**). In whole colon mucus extracts from specific pathogen free (SPF) mice colonized with the mouse G1 isolate *Lachnospiraceae* bacterium A4, we also detected A4 Fla1, Fla2, Fla3, and Fla4 but not A4 FlaB (**Table S12**).

To further characterize the flagellins that were detected, we conducted a flagellin phylogenetic analysis and included for comparison, any FlaB-like flagellin encoded by the individual isolates. The FlaB-like flagellins clustered separately (Clade 1) from the highly detected G1 flagellins which formed 2 distinct clusters (clades 2 and 3) with *Eubacterium* sp. 14-2 and *Roseburia inulinivorans* having flagellins represented in both (**Fig. S4D**). In contrast, all G2 flagellins detected formed a separate robustly supported cluster that we referred to as Clade 4 (**Fig. S4D**). A more expansive phylogenic analysis of all flagellins encoded by all organisms involved in the study, independent of their detection levels, recapitulated the clustering pattern of G1 and G2 flagellins with G1-encoded FlaB-like flagellins forming a separate highly supported cluster (**Fig. S4E**). Collectively, these data indicate that flagellins expressed by the genomes we classified as G1 or G2 are highly distinct, with G1 flagellins displaying a level of diversity consistent with the higher per-residue amino acid substitution rates illustrated in figure 1. In addition, the data argue that although 95% of the bacteria we classified as G1 encode a FlaB-like flagellin, these flagellins are not detected, and perhaps not expressed, at levels consistent with being packaged into flagella filaments. This conclusion was also supported by a phylogenic analysis of the 12 flagellins previously isolated from healthy human stool (*8*) which showed that 9 of the 12 clustered with reference G1 flagellins and separately from FlaB-like flagellins encoded by the respective genomes (**Fig. S5, A-B**).

To test the applicability of the foregoing characterization to the entire set of representative genomes, we generated a more comprehensive maximum likelihood tree that included all 518 flagellins encoded across the 163 genomes. The majority of flagellins, including each one analyzed above, segregated into the same 4 major clades. Smaller numbers of flagellins segregated into 5 subclades with 7 flagellins that did not fall into any of these clusters (**Fig. 4A**). This analysis further supported the separation of G1-encoded flagellins into the same 3 primary clades (clade 1-3) with only minor representation in other clades or subclades. Furthermore, the homogeneity of G2-encoded flagellins was again evident with over 80% of G2-encoded flagellins clustering in Clade 4 or subclade E and almost no representation in the G1-dominated clades (**Figure 4B**). In fact, for the lone G2 genome for which there is overlap (genome #128, *Eubacterium cellulosolvens*), 1 of 11 flagellins clustered in Clade 3 while the other 10 all clustered in Clade 4. Notably, among genomes #60 – #95, which did not meet our criteria for classification as G1 or G2, over 80% of genomes had at least one flagellin that clustered in clade 2 but approximately 50% of all flagellins encoded by these collective genomes were randomly associated with multiple other clusters (**Fig. 4B**). Altogether, these data support the generally divergent flagellin gene sequences of the bacteria we classified as G1 and G2 and validate the level of diversity within G1 and between the 2 groups.

**Fig. 4.**
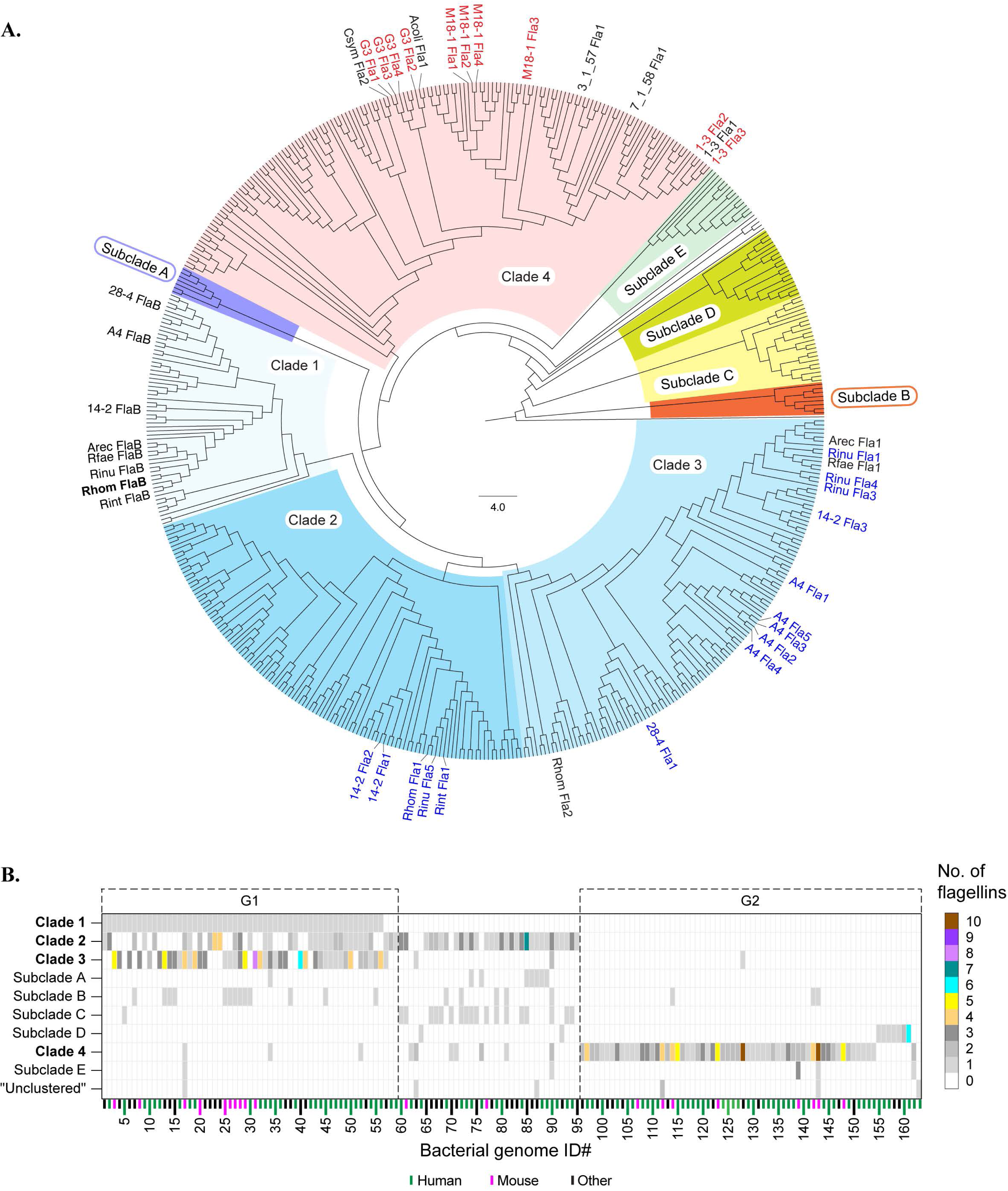
Phylogenic analysis of all flagellin genes encoded in the 163 representative genomes in the study. (**A**) Midpoint-rooted cladogram based on unbiased phylogenetic analysis of the 518 flagellin genes encoded by the 163 genomes. For reference, the positions on the tree of all flagellins detected in this study and the FlaB-like flagellins encoded by bacteria used in the study are indicated. Clusters in which at least 25% of the genomes were represented were considered a clade while others were considered subclades. Flagellins detected by mass spectrometry are highlighted (G1: blue, G2: red). (**B**) Heatmap based on (A) indicating the number of flagellins from each of the 163 genomes (x-axis) that are represented in each cluster. Dashed line indicates the use of the overlap between *flhA* to *flgF* distance and flagellin clustering to separate genomes into groups G1 and G2. [Acoli: *Anaerotruncus colihominis*, Arec: *Agathobacter rectale*, Csym: *Clostridium symbiosum*, Rfae: *Roseburia faecis*, Rint: *Roseburia intestinalis*, Rinu: *Roseburia inulinivorans*, 3_1_57: *Lachnospiraceae* bacterium 3_1_57FAA_CT1, 7_1_58: *Lachnospiraceae* bacterium 7_1_58FAA].

### G2 flagellins potently activate TLR5

Based on the flagellin expression patterns detected above, we focused exclusively on clade 2 or 3 flagellins of G1 bacteria G1 and the Clade 4 flagellins expressed by G2 as the premier mediators of any flagellin-dependent functions of the respective groups of commensals. We generated recombinant versions of the individual flagellins expressed by the *Lachnospiraceae* bacterium A4 (G1) and *Lachnospiraceae* M18-1 (G2) and used titrated doses of each to stimulate HEK-blue mTLR5 cells engineered to sensitively report activation of TLR5. With EC50 values of < 0.1 nM, all M18-1 flagellins more potently activated TLR5 than all A4 flagellins, with M18-1 Fla1 stimulating even more strongly than *Salmonella typhimurium* FliC (**Fig. S6, A-B**). Considering the reported expression of TLR5 in the murine proximal colon (*28*), we then generated organoid-derived epithelial monolayers from the proximal colons of WT mice. Stimulation of WT monolayers with flagellin-dominated protein extracts from *Lachnospiraceae* A4 failed to induce significant expression of Lcn2 or Cxcl1, even at high doses, whereas similar doses of M18-1 stimulated high expression of both Lcn2 and Cxcl1 (**Fig S6, C-D**). Therefore, despite slight variability in TLR5 activation, relative to G2 flagellins, the net effect of the entire pool of G1 flagellins is hypoactivation of TLR5-dependent effector pathways in epithelial cells.

We then tested the ability of aforementioned and additional representative recombinant flagellins shown to be expressed by mouse and human G1 and G2 isolates to directly activate intestinal epithelial cells. The majority of G1-derived flagellins tested were relatively weak stimulators of epithelial production of both Lcn2 and Cxcl1with average EC50 vales of >1nM in both cases (**Fig. 5A**). In contrast, the majority of G2-derived flagellins tested were stronger stimulators of both Lcn2 and Cxcl1, and in most cases the EC50 values were at least 10-fold lower than those of G1 flagellins (**Fig. 5A**). When we stimulated TLR5-sufficient (*Tlr5*+/+) or TLR5-deficient (*Tlr5*−/−) epithelial monolayers with single doses of the same G1- and G2-derived flagellins, the induction of Cxcl1 was completely ablated in TLR5-deficient epithelial cells (**Fig. S7A**), supporting an essential role for TLR5 in this cascade. We also injected SPF mice intraperitoneally with representative recombinant versions of flagellins confirmed to be expressed by G1 and G2 bacterial isolates. Similar to recipients of PBS-treated controls, we did not detect Cxcl1 and found minimal amounts of circulating Lcn2 in the recipients of G1-derived flagellins. In contrast, G2-derived flagellins, similar to the known pathogenic *Salmonella typhimurium* FliC, induced robust levels of circulating Cxcl1 and Lcn2 (**Fig 5B**), supporting the differential stimulatory capacity of G1- and G2-derived flagellins *in vivo*. Importantly, stimulation of lymphocyte-depleted peritoneal cavity cells from mice in which *Tlr5* was deleted in myeloid cells (*Lyz2*^Cre^.*Tlr5*^FL/FL^ *Tlr5*cKO mice), resulted in complete abrogation of Cxcl1 production, indicating that this response is largely dependent on myeloid cell TLR5 (**Fig. S7B**). We validated this effect *in vivo* using intraperitoneal injection of *Anaerotruncus* G3 Fla3 into wild type (*Tlr5*^FL/FL^) and *Lyz2*^Cre^.*Tlr5*^FL/FL^ littermates and detected robust circulating levels of Cxcl1 in wild type but not in *Lyz2*^Cre^.*Tlr5*^FL/FL^ mice (**Fig. S7C**). Together, these results demonstrate that G2- but not G1-encoded flagellins are strong stimulators of TLR5 on both epithelial and myeloid cells, and more generally, argue that intraperitoneal injection of flagellins induces Cxcl1 via TLR5 expressed by myeloid cells, and likely not via intracellular activation of NLRC4 as previously suggested (*8*).

**Fig. 5.**
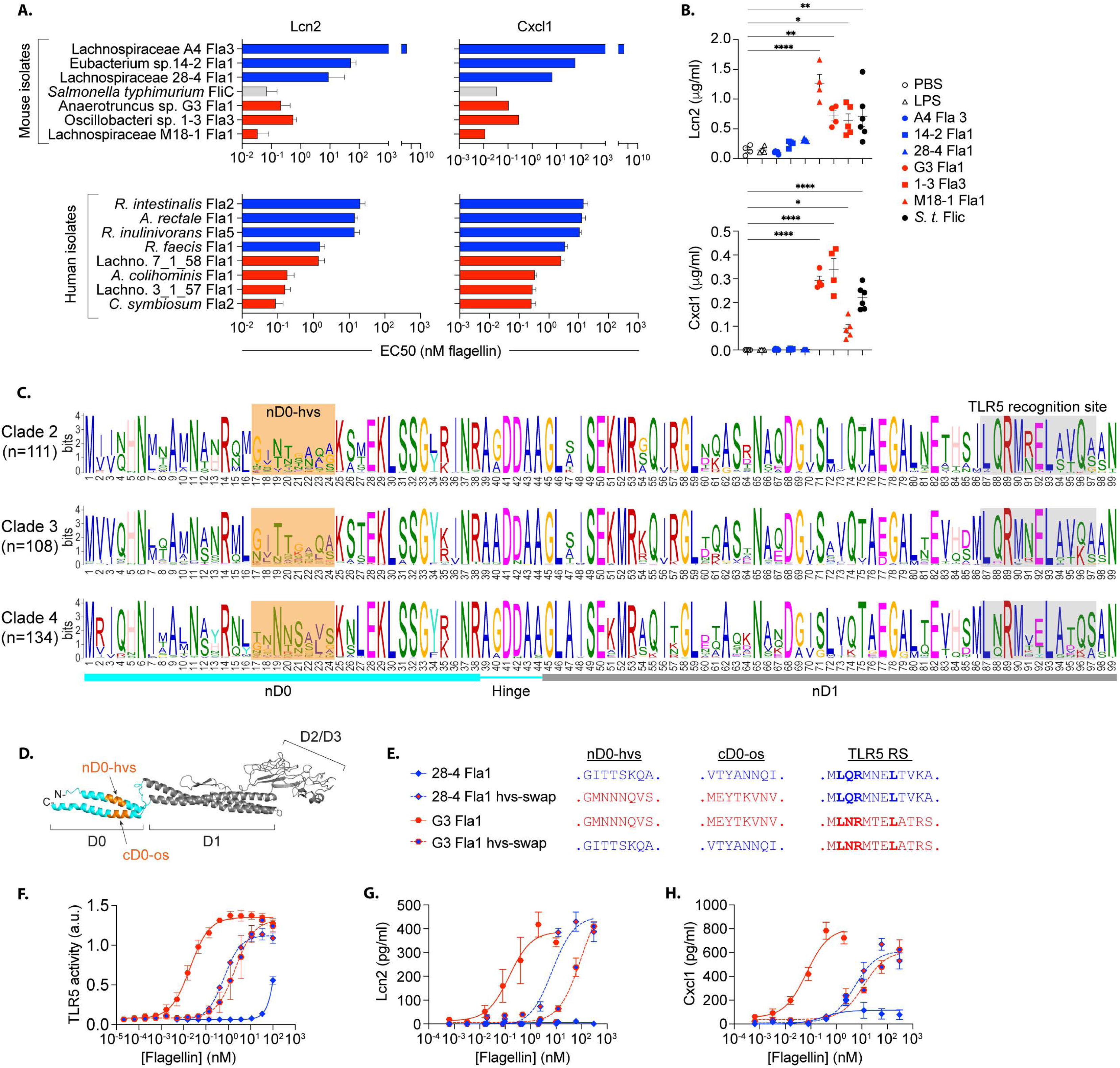
Distinct TLR5 stimulatory capacity of G1- and G2-encoded flagellins mediated by a non-conserved region of the D0 domain. (**A**) Mouse colon organoid-derived epithelial monolayers were activated with titrated doses of recombinant flagellins of mouse (upper) and human (lower) G1 and G2 isolates. Graphs show the average concentration of each flagellin required to produce half-maximal induction of Lcn2 and Cxcl1 (EC50, determined by non-linear regression analysis) after 24-hour stimulation. (**B**) Graphs show serum levels of Lcn2 and Cxcl1 2 hours after intra-peritoneal injection of the indicated recombinant flagellins (1μM in PBS) into naïve SPF mice. One group of mice received PBS as a negative control, and another received high dose LPS to ascertain that outcomes were not a result of endotoxin contamination of flagellin preparations. (**C**) Per-residue information in bits for sets of D0/D1 domain sequences for Clade 2 (upper), Clade 3 (middle), and Clade 4 (lower) computed using MEME Suite. Numbers of flagellins used in each computation are shown in parentheses. Location of the hypervariable region (nD0 hvs) is indicated by the orange shading. (**D**) Pictorial representation of flagellin basic structure highlighting the D0 domain (aqua) and 8 aa D0 epitopes (orange). (**E**) Sequences of the N- and C-terminal D0 epitopes as well as the TLR5 recognition sites of the native and hvs-swapped *Lachnospiraceae* 28-4 Fla1 and *Anaerotruncus* G3 Fla1. (**F**) Dose response curves showing TLR5 activity following 16-hour stimulation of HEK-blue mTLR5 cells with the respective native and hvs-swapped flagellins. Mouse colon organoid derived epithelial monolayers were also activated with titrated doses of the respective flagellins and the production of (**G**) Lcn2 and (**H**) Cxcl1 were determined by ELISA.

### The stimulatory potential of G1- and G2-derived flagellins is regulated by a defined region of the flagellin D0 domain

Flagellin monomers are comprised of relatively conserved D0 and D1 domains internal to the flagellar filament as well as hypervariable, external-facing D2 and D3 domains. The TLR5 recognition site is part of the D1 domain, but there is evidence that the D0 domain contributes to activation of TLR5 (*8, 29, 30*). We reasoned that sequences in the G1 and G2 D0 domains that contribute to this difference are likely to be highly distinct and can be used to guide initial interrogation of the factors that underpin the differential stimulatory capacity of G1 and G2 flagellins. Clustal-based alignment and peptide sequence analysis of the D0 and D1 domains of all flagellins considered in this study identified a highly variable 8 amino acid sequence in the N-terminal D0 (nD0) of flagellins in Clades 2, 3, and 4 (**Fig. 5C**), that we referred to as the nD0 hypervariable sequence (nD0-hvs). We used this finding to approximate the region of the entire D0 to be targeted, despite not identifying the same level of diversity in the C-terminal D0 (cD0) domain of flagellins in Clades 2, 3, or 4 (**Fig. S8A**). Interestingly that region of the nD0 is also highly diverse in FlaB-like flagellins, although notably distinct from that of other G1-derived flagellins (**Fig. S8B**). Considering previous findings implicating the cD0 in regulating flagellin stimulatory potential (*8, 29, 30*), and to preserve the structural integrity of that region of the D0 domains, we generated recombinant flagellins in which both the nD0-hvs and the opposing sequence on the cD0 (cD0-os) of the G2 flagellin *Anaerotruncus* G3 Fla1 were swapped for the corresponding sequences of the G1 flagellin *Lachnospiraceae* 28-4 Fla1 (G3 Fla1 hvs-swap) (**Fig. 5 D-E**). In all cases, the native TLR5 recognition site was left intact. When used to stimulate HEK-blue mTLR5 cells, the TLR5 stimulatory capacity of *Lachnospiraceae* 28-4 Fla1 hvs-swap was significantly increased relative to the native *Lachnospiraceae* 28-4 Fla1 with an approximately 2 log-fold reduction in EC50 (**Fig. 5F**). Conversely, The TLR5 stimulatory capacity of the *Anaerotruncus* G3 Fla1 hvs-swap was significantly less than parent *Anaerotruncus* G3 Fla1, with a greater than 100-fold increase in EC50 (**Fig. 5F**). An analogous trend was observed using reciprocal hvs-swaps featuring *Lachnospiraceae* A4 Fla3 and *Lachnospiraceae* M18-1 Fla1 (**Fig. S9, A-B**) or *Lachnospiraceae* 14-2 Fla1 and *Oscillobacter* sp. 1-3 Fla1 (**Figure S9, C-D**). Similarly, when organoid-derived epithelial monolayers were stimulated with titrated doses of the native and modified flagellins, the *Anaerotruncus* G3 Fla1 hvs-swap stimulated production of Lcn2 and Cxcl1 much less potently that than the native *Anaerotruncus* G3 Fla1, with EC50 values increased >1000 fold in each case. Conversely, the *Lachnospiraceae* 28-4 Fla1 hvs-swap more potently induced expression of Lcn2 and Cxcl1 than the native *Lachnospiraceae* 28-4 Fla1 with EC50 values that were reduced at least 1000-fold (**Fig. 5, G-H**). These results demonstrate that there exists at least one region the D0 domain of G1 and G2 flagellins that directly affects the intensity of TLR5 activation, the magnitude of downstream responses in epithelial cells, and possibly the pro-inflammatory potential of specific flagellated commensals.

### Distinct pattern of G1 and G2 bacterial abundance in health versus IBD

Previous studies showed reduced relative abundance of Eubacteriales, which at the time included the *Lachnospiraceae* family, within the fecal microbiota of CD patients relative to healthy individuals (*11–13*). This potentially diminishes a major immunoregulatory axis thereby favoring the pro-inflammatory milieu that is characteristic of IBD. We sought to determine whether this phenomenon applies to G1, G2, or both. Because the abundance of bacteria in the feces is not always reflective of events in the inflamed mucosa, we examined published 16S rRNA sequencing data from ileal biopsies of non-inflamed (NI) as well as CD and UC lesions (*31–33*). Briefly, we generated a BLAST database using ileal bacterial 16S rDNA sequences of patients deposited into the NCBI sequence read archives by Li et al., and Zhang et al. (*31, 32*) (Dataset 1) and Lloyd-Price et al., (*33*) (Dataset 2) (**Fig. 6A**). This database was then queried with human-derived Clostridia 16S rDNA sequences from our study to determine the relative abundance of taxa we have classified as G1 or G2. We established homology cutoffs of 99% for Dataset 1 which sequenced the 16S V3-V5 region, and 100% for dataset 2 which sequenced the V4 region (**Fig. 6A**). As internal controls, we examined the abundance of *Escherichia coli* (*E. coli*) and *Anaerostipes hadrus* (*A. hadrus*) which are known to be increased and decreased respectively in IBD (*34–36*). Overall, we detected 9 matches with bacteria we classified as G1 and 7 matches with G2. Of these, 2 G1 and 4 G2 represent independently correlated observations (**Fig. 6B**) but because of the 16S rDNA similarity between certain closely related species, greater than 75% correlation was observed with more than one species. In this case, we stringently advanced only one species for further consideration, although it is entirely possible that all species were represented in the dataset. Importantly, in every case, the overlap was restricted to bacteria belonging to the same group. Therefore, for bacteria that were present in a majority of individuals screened, we determined the relative abundance bacteria in CD and UC patients relative to NI controls. In CD biopsies, the average abundance of G1 bacteria *Simiaoa sunii*, *R. hominis*, and *R. inulinivorans* showed reduced median abundance, with *E. rectale* showing a statistically significant reduction in abundance (**Fig. 6C**). In contrast, the G2 bacteria *Clostridium symbiosum*, *Enterocloster clostridioformis*, and *Hungatella hominis* all showed statistically significant increases in abundance in CD biopsies relative to NI (**Fig. 6D**). Collectively, our results argue that organisms we classify as G2, which are arguably more potent drivers of inflammation, are expanded in CD lesions relative to healthy mucosa.

**Fig. 6.**
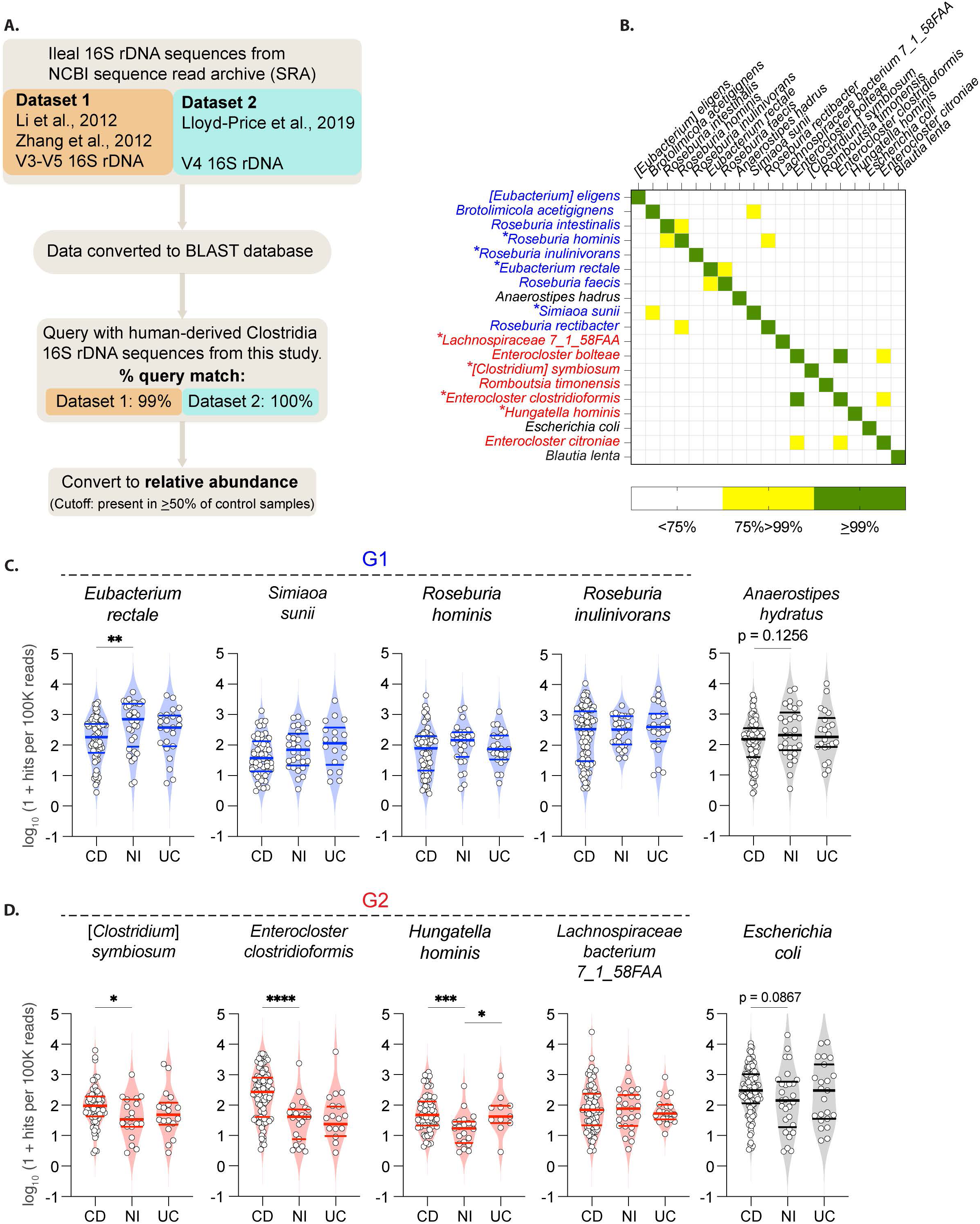
G1 and G2 bacteria are differentially abundant in healthy versus inflamed ileal biopsies. (A) Overview of the selection and querying of 16S sequences obtained from human ileal biopsies to determine relative abundance of G1 and G2 bacteria. (**B**) Heatmap showing correlation between the number of 16S sequence hits per 100,000 reads per individual for each human-derived genome in this study. For correlations of greater than 75%, only one representative genome was advanced for consideration. On the x-axis, bacteria classified as G1 or G2 are indicated by blue or red text, respectively, and black text indicates control bacteria known to be increased or decreased in IBD. Violin plots showing distribution of abundances (log10 hits per 100K reads) of (**C**) G1 and (**D**) G2 bacteria in ileal biopsies of Crohn’s disease (CD), and ulcerative colitis lesions. Each was compared to data from non-inflamed (NI) biopsies using Mann-Whitney test. Thick middle lines indicate medians, outer lines represent quartiles. *p<0.0332, **p<0.0021, ***p<0.0002, ****p<0.0001.

## DISCUSSION

In this study, we demonstrate that within the class Clostridia, flagellated bacteria that colonize the mammalian intestine can be segregated into at least 2 distinct consortia based on the diversity of flagellins and organization of motility genes present in their genomes. By every criterion used, the classification of G1 and G2 appears applicable to bacteria that have evolved with mice or humans. Evolutionary modeling predicts that host-associated bacteria that retain their flagella do so in conjunction with co-evolution of host control mechanisms particularly via TLR5 (*37*). Thus, our findings argue that mammalian hosts have facilitated the persistence of these taxonomically similar inhabitants with distinct pro-inflammatory potential via physical separation and immune-mediated tolerance. Overall, our study is a critical first step in elucidating the colitogenic potential of mucus associated commensal bacteria that possess the motility apparatus that can enable approach of, and association with, the intestinal epithelium.

The G1 organisms include notable members of the genus *Roseburia* that have been widely studied and were previously shown to promote tolerogenic responses *in vivo* (*9, 10*), as well as mouse isolate *Lachnospiraceae* A4, known to express flagellins recognized by circulating IgG in CD patients (*38*). The bacteria we define as G1 or G2 are also well tolerated during homeostasis, via mechanisms that involve RORγt+ Treg cells and anti-bacteria IgA. However, unlike G1, G2 bacteria promote robust pro-inflammatory responses in mice following mucus barrier disruption. Indeed, the 3 G1 and 3 G2 bacteria we have studied *in vivo* represent only a minor fraction of all that we have classified as G1 or G2. Although the mouse G2 isolates including those used in our gnotobiotic studies have not been widely studied in this context, this group also includes human-derived microbes determined to be present in IBD lesions in this and other studies, and bacteria such as *Hungatella hathewayi* and [*Clostridium*] *symbiosum* that have been positively correlated with IBD and colorectal cancer, respectively (*33, 39*). Ultimately, our study establishes a framework, and stratifies bacterial candidates, for continued exploration of the host-microbiota crosstalk occurring at the mucosal border during health and in development and chronicity of gut inflammation and downstream comorbidities.

Despite the flagellin gene diversity apparent in the genomes of G1 organisms, our results align with previous findings (*6*) demonstrating that that only some flagellins are expressed at levels indicative of packaging into flagella filaments and are thus essential for motility. Our findings argue that flagellins which cluster in what we refer to as Clades 2 and 3 are the primary G1-encoded flagellins expressed and sensed by host epithelial TLR5 under homeostatic conditions, where the sub-optimal activation enables tolerance of G1 bacteria. It is likely that multiple microbial products of G2 bacteria may trigger epithelial activation at homeostasis and especially following a mucus disrupting insult. However, our findings position the G2-encoded flagellins as possible mediators of the pro-inflammatory potential of these bacteria, likely via the interaction with epithelial TLR5. Importantly, our data indicate that the G1- and G2-encoded flagellins also differentially stimulate TLR5 expressed on myeloid cells, increasing the potential for sustained G2-dependent inflammation following barrier disruption.

Our comparisons of the D0 domains of all G1 or G2 flagellins identified a distinct 8-amino acid region of the nD0 domain, but interestingly, not the cD0 domain, although this nD0 hypervariable sequence is adjacent to a region of the cD0 domain previously shown to modulate TLR5 activation. Our findings are consistent with previous findings that structural features, distinct from the known TLR5 recognition site, significantly affect the stimulatory potential of flagellins (*8, 29, 30*). The size and diversity of the D0-hvs, coupled with the variability in TLR5 activation strength within and across groups may enable even more fine-tuning of these interactions providing an added layer of commensal-dependent regulation of host responses.

A previous study utilized the *R. hominis* FlaB to demonstrate the concept of “silent flagellin recognition” of human TLR5, relative to flagellin from pathogenic *Salmonell*a (*8*). Interestingly, despite *R. hominis* being a human isolate, results obtained following injection of *R. hominis* FlaB into wild type and *Tlr5*−/−.*Nlrc4*−/− mice to assess *in vivo* immune responses to this representative silent flagellin led to the conclusion that “silent recognition” of flagellin by TLR5 is not species-specific. Consistent with this, we found that G1 or G2 flagellins clustered together independent of the host from which the bacteria were previously obtained, and our *in vitro* experiments showed similar patterns of TLR5 stimulation by recombinant flagellins whether encoded by mouse- or human-derived isolates. Surprisingly, however, in cultures of both mouse and human bacteria isolates including *R. hominis*, we failed to detect significant levels of *R. hominis* FlaB or similar flagellins encoded by the mouse- or human-derived G1 bacteria tested. We also failed to detect *Lachnospiraceae* A4 FlaB among the flagellins found in A4-colonized SPF mice, and flagellins previously isolated from human stool (*8*) were distinct from the “FlaB-like” flagellins encoded by the respective genomes. The reason for this is currently unclear but it is worth noting that experiments demonstrating TLR5 hypoactivation by recombinant *R. hominis* FlaB and similar flagellins utilized only recombinant flagellins, and actual expression either *in vitro* or *in vivo* was never demonstrated (*8*). The very low-level, if any, detection of FlaB flagellins in our study, the inability to identify FlaBs in human stool, and the belief that *R. hominis* FlaB activates the NLRC4 inflammasome (*8*), suggest a mechanism of action distinct from that identified for the G1 and G2 flagellins studied here, which seems to be strictly dependent on TLR5.

Anti-flagellin IgG and CD4 T cell reactivity are believed to implicate specific bacteria in Crohn’s disease (CD) (*14, 40, 41*), and seroreactivity to FlaX and *Lachnospiraceae* A4 Fla2 can precede development and diagnosis of CD by up to a 5 years (*42*), raising the potential for informing preventative interventions in susceptible individuals. We recently showed that the same anti-flagellin reactivity is present in infants from multiple geographic regions, suggestive of tolerogenic responses that help facilitate early life colonization by flagellated bacteria (*43*). Thus, despite strong clinical data, it is still unclear whether anti-flagellin antibodies in Crohn’s disease are a cause or consequence of colonic inflammation (*44*). We recently mapped the major epitope of *Lachnospiraceae* flagellins recognized by anti-flagellin IgG and found that these antibodies primarily bind the hinge region linking the D0 and D1 domains (*43*), which we now know based on the current study, is highly conserved across flagellins in Clades 2, 3, and 4, which encompass the majority of G1 and G2 flagellins. Thus, the actual target(s) associated with the anti-A4 Fla2 responses detectable in Crohn’s disease could be flagellin(s) derived from G2 bacteria or any number of G1 bacteria. At a minimum, these observations combine to argue that anti-flagellin antibodies may not necessarily track with the colitogenicity of any bacterium or the pro-inflammatory potential of the flagellins it encodes.

Despite the widespread association of anti-flagellin reactivity with Crohn’s susceptibility and severity, studies have shown that bacteria in the family *Lachnospiraceae*, believed to be the primary producers of these flagellins, are actually reduced in IBD patients (*11–13*). Therefore, despite the caveat that it is difficult to get species level resolution of microbial signatures from bacterial 16S rDNA sequencing, we attempted to mine publicly available data generated by sequencing of mucosal biopsies of IBD patients and healthy subjects, to determine whether or not the bacteria we classified as G1 versus G2 are detectable and at least semi-quantifiable in Crohn’s disease lesions, instead of less informative fecal samples. Importantly, we could identify several sequences that uniquely mapped to human G1 and G2 isolates. In general, the G1 sequences identified in healthy controls showed slight but mostly non-statistically significant reductions in abundance in CD lesions. In contrast, 3 of 4 G2 bacteria identified showed statistically significant increases relative to controls. Thus, in addition to the mouse G2 isolates used in our gnotobiotic experiments being more pro-inflammatory, the human candidates identified in the published data also appear to be enriched in CD biopsies. At a minimum, our findings argue that *Lachnospiraceae* bacteria are likely not uniformly depleted during chronic gut inflammation, and caution against broad generalizations regarding microbial dynamics in IBD based on fecal abundance at higher taxonomic levels. More importantly, our results point to specific members of the class Clostridia within the healthy human microbiota as potential contributors to CD-associated inflammation perhaps due to the ability to withstand inflammatory conditions and induce inflammatory pathways in gut epithelial cells.

### Study limitations

The inflammation observed in G2-colonized mice given low dose DSS suggest that these bacteria are potentially colitogenic in this model. Although recombinant versions of flagellins encoded by G2 bacteria induced inflammatory gene expression in colonic epithelial monolayers, is still not known whether the colitis observed is flagellin-dependent, and if so, whether these outcomes are also impacted by modification of the nD0, cD0, or both. Further work will be necessary, particularly for the human isolates in combination with human TLR5, to determine whether the phenomena described herein are replicated in the presence of other G1 or G2 bacteria, and the extent to which the apparent pro-inflammatory functions of G2 bacteria are dependent on TLR5 and/or flagellins. Considering that anti-flagellin responses have been implicated in Crohn’s disease, it would be valuable to examine the described phenomena in a model that more closely resembles Crohn’s disease and to better understand flagellin expression patterns *in vivo* during both homeostasis and inflammation, and to determine whether the stimulatory patterns observed on mouse TLR5 are also seen on human TLR5. Finally, the effect of the D0 domain described by us and others, or overall differences between G1 and G2-encoded flagellins, could be linked to structural features of both, but flagellin structures were not explored in this study.

## MATERIALS AND METHODS

### Study Design

The objective of this study was to determine whether differences in motility apparatus of colon mucus associated commensal bacteria might provide a basis for exploring the functional diversity of these organisms, including potential flagellin-dependent pro-inflammatory capabilities that may underlay the colitogenicity of specific bacteria. We focused initially on members of the class Clostridia within the phylum Bacillota and identified 163 representative flagellin-encoding genomes of bacteria previously isolated from the guts of humans, mice, and other hosts. After observing that the arrangement of specific motility loci and the diversity of encoded flagellins can indeed distinguish many of the bacteria, we used gnotobiotic approaches to explore the homeostatic and pro-inflammatory functions of representatives of 2 clearly demarcated groups *in vivo* and used *in vitro* assays to determine the TLR5 activating capacity of the flagellins they detectably express. This led us to the identification of a single motif in the D0 domain of the flagellins studied affects the differential stimulatory capacity of the specific flagellins. Finally, we analyzed publicly available data to determine the relative abundance of bacteria classified into the 2 groups in biopsies from non-inflamed versus Crohn’s disease and ulcerative colitis lesions.

### Mice

All mice used in this study were on the C57BL/6 background and were used in accordance with the UAB Institutional Animal Care and Use Committee guidelines. Previously reported IL-10 reporter transgenic (10BiT) mice (*19*) were used under both SPF and germ-free (GF) conditions. *Lyz2*^Cre^.*Tlr5*^FL/FL^ mice were generously provided by Dr. Stavros Garantziotis of the National Institute of Environmental Health Sciences (NIEHS). To generate gnotobiotic mice, adult GF 10BiT mice were gavaged with pure cultures of bacteria and colonization was confirmed by PCR after 14 days. Colonized females were mated with GF males to generate progeny for use in the experiments. The gnotobiotic colonies were propagated by mating the progeny of the originally colonized animals. Gnotobiotic mice were screened for the presence of all the respective G1 and G2 strains at 6 weeks of age, prior to being used in experiments.

### Genome identification

Gut derived genomes were identified using the metadata for the genomes provided by NCBI. For genomes missing this data we searched the literature for papers describing the isolation of the particular strain. Mapping of motility genes was done using a mixture of genome and protein analysis tools and manually assembled. Bacterial genomes were aligned in each motility locus using the PATRIC “Compare Region Viewer”. Similarly annotated genes that aligned in each genome using a published approach (Neville et al., 2013) as a guide were collated in excel to create loci maps. Unannotated genes from loci maps of each genome were identified using the translated protein sequence in UniProt domain analysis. Proteins that had domain matches corresponding to domains from annotated proteins encoded in the same position in the locus were identified in the locus map as that protein.

### Calculation of motility gene distance

Using Neville et. al. (*6*) as a guide, the location of annotated motility genes from the *flgB*-*fliA* & *mbl*-*flgJ* loci was mapped as PATRIC (*45*) protein encoding gene (PEG) numbers for all the genomes studied. The *flhA* gene within the *flgB*-*fliA* locus showed minimal divergence across genomes and therefore the *flhA* PEG number was used as a proxy for relative location of the *flgB*-*fliA* locus of each genome. We then noted PEG numbers of the *flgF* from each genome. The difference between these two PEG numbers represents the gene distance shown in Figure 1B. We confirmed that all the genomes with a large gene distance also contained a complete *mbl*-*flgJ* locus.

### Identification of flagellin genes

Flagellins were identified using the annotation information provided with each genome. We also did a blast of all 163 genomes using *Roseburia hominis* A2-183 Fla1 and *Lachnospiraceae* bacterium 3_1_57FAA_CT1 Fla1 as queries to confirm we were not missing any flagellins. All flagellin genes were identified by the annotation information with 3 exceptions, 2 of which fall on the broken edge of contigs and thus escape annotation. *Acetivibrio ethanolgignens* strain ACET-33324 Fla was identified using BLAST, and *Eubacterium* 14-2 Fla1 and Fla2 which were previously cloned by our group (Duck et al., 2007). We also provided the DNA that was used for the genome sequencing of *Eubacterium* 14-2 currently in NCBI.

### Determination of flagellin gene diversity within individual genomes and across clusters of genomes

Flagellins were analyzed using molecular evolutionary genetics analysis (MEGA) (*46*) The sequences were aligned using Multiple Sequence Comparison by Log-Expectation (MUSCLE) (*47*) within MEGA. Aligned flagellins were grouped either by genome (163 groups) or by genome cluster (3 groups). Estimates of average evolutionary divergence over sequence pairs within groups was conducted for both the 163 individual genomes or each of the 3 genome clusters as a group. Amino acid positions with less than 95% site coverage were eliminated, meaning that fewer than 5% alignment gaps, missing data, and ambiguous bases were allowed at any position. Standard error estimates were conducted using the bootstrap procedure (1000 replicates). Analyses were conducted using the JTT matrix-based model (*48*). The rapid generation of mutation data matrices from protein sequences. Computer Applications in the Biosciences 8: 275-282.

### Phylogenetic Analysis

Flagellins were analyzed using molecular evolutionary genetics analysis (MEGA) (*46*). Amino acid positions with less than 95% site coverage were eliminated, meaning that fewer than 5% alignment gaps, missing data, and ambiguous bases were allowed at any position. The sequences were aligned using Multiple Sequence Comparison by Log-Expectation (MUSCLE) (*47*). 518 aligned Clostridia sequences were then subjected to model analysis to determine which Maximum Likelihood substitution model best fit the data (**Data File S1**). The same model was used for all phylogenetic tree analyses. The flagellin phylogenetic association was inferred using the Maximum Likelihood method and LG (Le_Gascuel_2008) model (*49*), and we selected the tree with the highest log likelihood. To obtain an initial tree for the heuristic search, Neighbor-Joining (NJ) and BIONJ (*50, 51*) algorithms were applied to a matrix of pairwise distance estimates obtained using the JTT model (*48*) and then selecting the topology with superior log likelihood value. FigTree software (*52*) was used to present the tree as a radial, mid-point rooted cladogram. The tree in Figure S4D represents 31 amino acid sequences and 216 positions, Figure S4E represents 58 sequences and 200 positions, Figure S5B represents 58 sequences and 200 positions, and Figure 4A represents 519 sequences and 208 positions.

### Colon lymphocyte isolation

Colon was collected and cleared of visible luminal contents by flushing with cold PBS. The colon was cut open longitudinally then cut into 1cm pieces and collected into cold H2 media (HBSS with 2% fetal bovine serum). Tissues were gently rotated for 20 minutes in H2/DTT/EDTA buffer at 37°C. Media was decanted and tissue was briefly washed in H2. After decanting H2, tissue pieces were minced in a gentleMACS C Tubes (Miltenyi Biotec, 130-093-237) containing 1000U/mL of collagenase IV and 200 ug/mL DNase I. The tissue was then rotated at 37°C for 45 minutes. After incubation, the tissue was placed on the gentleMACS Dissociator (Miltenyi Biotec) and ran through the program m_intestine_01 a total of two times. Cells were then strained through a 100μM filter, pelleted, and resuspended in 37% percoll in R2 media (RPMI containing 2% FBS and 1% Penn/Strep). Cells were centrifuged for 20 minutes at room temperature with no brakes. Single cell suspension from the Percoll interface were collected and washed in cold R10 media (RPMI containing 10% FBS and 1% Penn/Strep) in preparation for further analysis.

### Microbial flow cytometry analysis (mFlow)

Mouse pellets were collected in 0.5ml of 1X protease inhibitor cocktail (Roche, cat. # 05892953001) and homogenized. Large debris was removed by centrifugation at 1,000 rpm for 1 min, and resulting supernatant was centrifuged at 6,000rpm for 5 min to pellet bacteria. The bacteria pellets were resuspended in 1ml of staining buffer (1%BSA) and passed through a 35μm nylon filter. The bacteria were washed twice with staining buffer and stained with SYTO BC nucleic acid dye (ThermoFisher Scientific S34855) at 1:4000 dilution and anti-mouse IgA-PE (clone mA-6E1, Invitrogen cat. # 12-4204-83) at 1:100 dilution for 20 min on ice. SYTO BC+ gates for total bacteria were set based on staining of germ-free pellets, and gates for IgA-bound bacteria were determined based on staining of pellets from IgA-deficient mice. Samples were acquired on a Northern Lights 3L cytometer (Cytek), and data were analyzed by FlowJo software.

### Cell staining, antibodies, and flow cytometry

Single cell suspensions were incubated on ice for 15 minutes with Zombie Near-IR Fixable dye (Biolegend, 423106) diluted 1:1250 in PBS. Cells were then washed and blocked with TruStain FcX PLUS (Biolegend, 156604) on ice for 15 minutes. The following antibodies were diluted in FACS buffer (PBS containing 2% FBS, 1% Penn/Strep and 2mM EDTA) with Brilliant Stain Buffer Plus (BD Horizon, 566385) and 50 μl of this cocktail was added to cells followed by incubation on ice for 25 minutes: CD90.1 Brilliant Violet 711 (Biolegend, 202539), CD4 APC (Biolegend, 100412), TCRβ PE/Cyanine7 (Biolegend, 109222), and CD45 Brilliant Violet 785 (Biolegend, 103149). Cells were then washed and permeabilized with the Fix/Perm buffer of the Foxp3/Transcription Factor Staining Buffer Set (Invitrogen, 00-5523-00) per the manufacturer’s instructions. The intracellular stain was performed at room temperature for 45 minutes using the following antibodies: Foxp3 Alexa Fluor 488 (Invitrogen, 53-5773-82), RORγt PE (BD Biosciences, 562607), and Helios PE/Dazzle 594 (Biolegend, 137232). Cells were acquired on a Northern Lights flow cytometer (Cytek Biosciences) and data were analyzed using FlowJo v10.10 software.

### Colon Tissue Staining

Whole colons with fecal contents were collected into Methacarn, Carnoy’s fixative (bioWORLD, 21730013-2) and incubated overnight. Samples were washed twice for 30 minutes with 100% methanol, then twice for 20 minutes with 100% ethanol. The colon was then cut into 1 cm proximal and distal sections and placed in histology cassettes in xylene. Tissue was embedded in paraffin and 5 μm sections were cut onto glass slides.

#### Fluorescence in-situ hybridization (FISH), immunofluorescence staining, and analysis

Slides were de-waxed at 56-60°C for 30 minutes and then placed in pre-warmed xylene for 15 minutes. The slides were placed in xylene an additional 2 times for 15 minutes each. Slides were then placed in 100% ethanol for 15 minutes followed by 95% ethanol for 15 minutes. The slides were placed in PBS and crosslinked with an ultraviolet light subtype-C for 20 minutes. Slides were air-dried and hybridized overnight at 50°C with 1 μg/ul of the universal bacterial 16S probe EUB 338 conjugated to Alexa 594 (Invitrogen). Slides were then stained with 5 μg/ml of wheat germ agglutinin (Alexa 488) for 1 hour at room temperature, followed by 1 μg/ml of DAPI for 5 minutes at room temperature. After all staining and washing steps were complete, coverslips were mounted using ProLong Gold antifade (Fisher, P36934). Slides were visualized using the Nikon A1R confocal microscope using the 60x oil-immersion objective.

#### H&E staining and analysis

Slides were stained with hematoxylin and eosin (H&E) for histologic assessment by a pathologist who was blinded to the experimental groups. Histological severity was determined based on a previously described scheme (*53, 54*) with modifications. Briefly, the follow parameters were graded on a scale from 0 (normal) to 4 (severe) to determine the epithelial score: extent of epithelial damage, epithelial hyperplasia, goblet cell depletion, epithelial depletion/necrosis, epithelial erosion/ulcer. The following parameters were graded on a scale from 0 (normal) to 4 (severe) to determine the inflammation score: extent of inflammation, depth of inflammatory infiltrate, crypt exudate, edema, and transmural inflammation. Total histology score was determined as the sum of epithelial and inflammatory scores. Representative images were collected using Nikon Eclipse Ci microscope and analyzed with NIS-elements software.

### Preparation of recombinant flagellins

Flagellin protein sequences were derived from published genomic data. Flagellins of interest were synthesized by GeneArt, codon optimized for expression in *E. coli* and cloned into expression vector pET100/D-TOPO. Plasmids were transformed into BL21 Star^TM^ (DE3). Protein production was induced with 0.5mM IPTG and cell pellets were extracted with 8M urea. Flagellins were purified using nickel column chromatography, washed, eluted, then dialyzed against 10mM Tris.

### Flagellin injections

Groups of 3-4 mice were each injected intraperitoneally with 1μM of flagellin in sterile PBS. Negative control mice were given an equal volume of PBS. At 2 hours post injection, serum was collected and spun at 16,000 x g for 10 minutes at 4°C. Serum was collected and frozen at −80°C until ready to analyze via ELISA.

### Enzyme Linked Immunosorbent Assay (ELISA)

Mouse CXCL1/KC DuoSet ELISA (R&D Systems, DY453) and Mouse Lipocalin-2/NGAL DuoSet ELISA (R&D Systems, DY1857) kits were utilized according to the manufacturer’s protocol. The TMB Substrate kit (Fisher, PI34021) was used to develop the assay, and results were read on a Biotek Synergy HTX plate reader. Wavelength correction of absorbance values was done via reading of the plate at OD540nM. Maxisorp ELISA plates (Fisher, 12-565-135) were used for all assays.

### HEK-Blue mTLR5 Assay

HEK-Blue mTLR5 cells were maintained and passaged, and the QUANTI-Blue assay was performed according to the manufacturer’s instructions (Invivogen). Briefly, 20μl of various concentrations of flagellin in PBS was placed in a 96-well, flat-bottom cell culture dish in duplicate. To each well, 20,000 freshly diluted HEK-Blue cells resuspended in 180μl of HEK-Blue Detection (SEAP) Medium were added. The plate was incubated at 37°C in 5% CO_2_ for 16 hours and read at 620nm in a Synergy HTX (BioTek) plate reader.

### Generation and stimulation of colonic organoid-derived epithelial monolayers

Mouse intestinal organoids were generated using a previously described protocol (*55*). Briefly, a 0.3 cm^2^ biopsy from mouse proximal colon tissue was taken and digested at 37°C with collagenase I solution (2mg/mL collagenase I and 50μg/mL gentamicin) in wash buffer (10% FBS, 1% Pen/Strep, 1% L-glutamine, 1% fungizone, and 0.1% gentamicin) until 50-80% of epithelial units were separated (approximately 30-50 minutes). Resulting cells were pelleted and washed, filtered through a 70μm cell strainer, washed, resuspended in cold Matrigel Matrix Basement Membrane (Corning, 354234), and plated at 15μl per well in a 24-well plate. Organoids were passaged as previously described (*55*) via manual scraping of the Matrigel dome in 500μL of 0.1% EDTA in PBS at least once before differentiation into epithelial monolayers.

To generate epithelial monolayers, organoids were trypsinized, filtered through a 40 μm strainer, washed and counted. Cells were plated in previously coated (1:30 Matrigel-PBS) 96 well plates at 100,000 cells per well in 200 μl of 5% L-WRN Conditioned Media prepared as previously described (*55*). On day 2, confluent monolayers were stimulated for 24 hours with recombinant flagellin. After stimulation, supernatant was collected and centrifuged and stored at −20°C.

### Statistical Analysis

To compare 2 or more independent groups, we performed ordinary 1-way ANOVA with Tukey’s multiple comparison test and two-tailed *P* values < 0.05 were considered significant. For multiple comparisons involving grouped results we performed ordinary 1-way, or 2-way analysis of variance (ANOVA) followed by Sidak’s multiple comparison test or Brown-Forsythe and Welch ANOVA with Dunnett’s T3 multiple comparisons test. All analyses and generation of graphs were done using PRISM software version 10 (GraphPad Software, Inc.). Error bars indicate standard error of the mean (SEM), and horizontal bars indicate means, medians, or quartiles as indicated in the figure legends.

## Supplementary Methods

### Bacterial strains and growth conditions

All strains used in this study were obtained from the biorepository of the UAB Microbiota Culture and Analytics Resource (MCAR). *Lachnospiraceae* bacterium A4, *Eubacterium* sp. 14-2, *Lachnospiraceae* bacterium 28-4, *Lachnospiraceae* bacterium M18-1, *Oscillobacter sp.* 1-3, and *Anaerotruncus sp.* G3 were previously isolated from mouse cecum (*16*). *Roseburia* intestinalis L1-82 (*56*), *Roseburia hominis* A2-183 and *Roseburia inulinivorans* DSM16841 were originally isolated from human feces (*57*) and samples were purchased from Leibniz Institute DSMZ. All bacterial strains were grown under anaerobic conditions using a Coy Lab vinyl anaerobic chamber under a 90% N_2_, 5% CO_2_, and 5% H_2_ gas mix. Bacteria were grown at 37°C on M2GSC media [10 g/L Bacto Casitone, 2.5 g/L Bacto yeast extract, 4 g/L NaHCO_3_, 2 g/L glucose, 2 g/L soluble starch, 2 g/L cellobiose, 30% rumen fluid, 15% mineral solution I (3 g/L K_2_HPO_4_), 15% mineral solution II (3 g/L KH_2_PO_4_, 6 g/L (NH_4_)_2_SO_4_, 6 g/L NaCl, 0.6 g/L MgSO_4_, 0.6 g/L CaCl_2_), 0.01% resazurin, and 1 g/L cysteine HCL] for 1-3 days.

### Generation of gnotobiotic cohorts

Germ-free (GF) 10BiT female mice were transplanted with either the G1 or G2 trio of organisms and colonization was confirmed 14 days later. Gnotobiotic colonies were established by mating these females with germ-free males. Fecal PCR at 6-8 weeks of age was used to confirm colonization by all bacteria. The gnotobiotic colony was maintained by breeding the offspring of the original and subsequent matings. Thus, all gnotobiotic mice examined in this study naturally acquired the bacteria via maternal transmission in early life.

### Quantification of bacterial colonization of intestinal mucus layer

Cecum and colon were cut open longitudinally. Visible luminal debris were completely removed by washing with cold PBS. The colon was then cut into cecum, proximal, middle, and distal sections. Mucus from each intestinal section was scraped into pre-weighed tubes and mucus weight was recorded. DNA was extracted using the Wizard Genomic DNA Purification Kit (Promega, A1120) with a modified protocol starting at Step 3 with a 1-hour incubation at 55°C. After extraction, qPCR was run using the SsoAdvanced Universal SYBR Green Supermix (BioRad, 1725275) and the qPCR specific primers (**Table S13**). The qPCR was run on a QuantStudio 3 thermocycler using the following parameters: 95°C for 10 minutes, 45 cycles of 95°C for 15 seconds and 61°C (for isolates 14-2, A4, M18-1, or G3) or 63°C (for isolates 28-4 or 1-3) for 30 seconds, then followed with a melt curve. To quantify the C_T_ values, we generated a standard curve utilizing a pMA-RQ plasmid containing our PCR product of interest synthesized by Thermo Fisher GeneArt. Copies of bacteria per μl were normalized to mucus weight. Data were analyzed using the online Design and Analysis App provided by Thermo Fisher Connect.

### Isolation and enrichment of bacterial proteins

Bacterial cultures were spun at 1800 x g for 10 minutes at room temperature. Bacterial pellets were washed with TE buffer (10 mM Tris-HCl, 0.1 mM EDTA) then resuspended in 12 mL of TE buffer. The suspension was poured into the stainless-steel mini container (Grainger: model# MC2, cat. # 45H25) attached to a Waring Commercial blender base (Waring Commercial: model# 7011BU). and run at high speed for 2 minutes. Blended bacteria were spun in an Optima ultracentrifuge at 45,000 rpm for 1 hour, the supernatant was discarded, and the pellet was resuspended in 500 μl of TE buffer. The pellet was then washed twice with TE buffer, spun at 10,000 g for 20 minutes and each time the supernatant was collected. Combined supernatants from each bacterial preparation containing extracellular protein extracts were quantified via BCA assay and run an SDS-PAGE on a 4-20% gradient Novex™ Tris-Glycine Mini Protein Gels (Invitrogen, XP04200BOX) that was later stained with Coomassie Blue to qualitatively confirm the presence of proteins in the prep.

### Proteomics Analysis

#### Sample Preparation

Proteomics analyses were carried out as previously referenced (*58*) with minor changes. All protein extracts were attained using M-PER™ Mammalian Protein Extraction Reagent (Thermo Fisher Scientific, Cat. # 78501) and quantified using Pierce BCA Protein Assay Kit (Thermo Fisher Scientific, Cat. # PI23225). As experimentally determined, 2-5ug of protein per sample were diluted to 35µL using NuPAGE LDS sample buffer (1x final conc., Invitrogen, Cat. # NP0007). Proteins were then reduced with DTT and denatured at 70°C for 10min prior to loading everything onto Novex NuPAGE 10% Bis-Tris Protein gels (Invitrogen, Cat. # NP0315BOX) and separated as a short stack (10 min at 200 constant V). The gels were stained overnight with Novex Colloidal Blue Staining kit (Invitrogen, Cat. # LC6025). Following de-staining, each lane was cut into a single MW fraction and equilibrated in 100 mM ammonium bicarbonate (AmBc), each gel plug was then digested overnight with Trypsin Gold, Mass Spectrometry Grade (Promega, Cat. # V5280) following manufacturer’s instruction. Peptide extracts were reconstituted in 0.1% Formic Acid/ ddH_2_O at 0.1µg/µL.

#### Mass Spectrometry

Peptide digests (8µL each) were injected onto a 1260 Infinity nHPLC stack (Agilent Technologies) and separated using a 75-micron I.D. x 15 cm pulled tip C-18 column (Jupiter C-18 300 Å, 5 microns, Phenomenex). All data were collected in CID mode. Following each parent ion scan (300-1200m/z @ 60k resolution), fragmentation data (MS2) were collected on the topmost intense 10 ions @7.5K resolution. For data dependent scans, charge state screening and dynamic exclusion were enabled with a repeat count of 2, repeat duration of 30s, and exclusion duration of 90s.

#### MS Data Conversion and Searches

The XCalibur RAW files were collected in profile mode, centroided and converted to MzXML using ReAdW v. 3.5.1. The mgf files were created using MzXML2Search (included in TPP v. 3.5) for all scans. The data were then searched using SEQUEST (Thermo Fisher Scientific). Searches were performed with a tailored species-specific subset of the UniProtKB database.

#### Peptide Filtering, Grouping, and Quantification

The list of peptide IDs generated based on SEQUEST search results were filtered using Scaffold (Protein Sciences, Portland Oregon). The filter cut-off values were set with minimum peptide length of >5 AA’s, with no MH+1 charge states, with peptide probabilities of >80% C.I., and with the number of peptides per protein ≥2. The protein probabilities were set to a >99.0% C.I., and an FDR<1.0%. Relative quantification across experiments was performed via spectral counting, and when relevant, spectral count abundances were then normalized between samples.

### RNA extraction and qPCR

PBS was added to cells and centrifuged at 16,000 x g for 7 minutes. The cell pellet was resuspended in 1 ml of trizol and disrupted by sonication. 200 μl of chloroform was added and mixture was shaken for 15 seconds followed by incubation at room temperature for 5 minutes. The mixture was centrifuged at 12,000 g for 15 minutes at 4°C and the upper colorless aqueous layer was collected in a new tube containing an equal volume of isopropanol. The mixture was inverted several times then placed at −20° C to precipitate the RNA overnight. The next day, the RNA was centrifuged at 12,000 g for 15 minutes at 4°C. The pellet was washed with 70% ethanol and vortexed then centrifuged again. The pellet was air dried for 5 minutes then resuspended in 100 μl of H_2_O and placed on a 60°C heat block for 10 minutes. Extracted RNA was reverse transcribed using the High-Capacity cDNA Reverse Transcription Kit (ThermoFisher, 4368814) according to instructions. qPCR was performed using the SYBR Green Mastermix. The sequences for the primers utilized are indicated in the **Table S13**. The fold change for each gene was calculated using the ΔΔCt method with the *Eef2* gene as the housekeeping gene.

### RNA-Seq Analysis

Adapters were trimmed from raw paired-end sequence data using trim_galore 0.4.4 (*59*). Reads were aligned using STAR 2.5.2 with default parameters and the Mus musculus GRCm39 reference (*60*). Aligned reads were counted using HTSeq-Count 0.6.1p1 with the stranded option set to “no” and using the GTF genome reference file GCF_000001635.27_GRCm39_genomic.gtf (*61*). Three replicates each of GF, G1 and G2 mice were analyzed for differential expression using the edgeR package (*62*). Only genes with counts per million greater than 1 for three or more samples were considered for further analysis. The contrasts G1 vs. GF and G2 vs. GF were evaluated to identify differentially expressed genes using a GLM approach in edgeR.

### Lcn2 ELISA

Fresh fecal pellets were collected on Day 7, weighed, and re-suspended in PBS containing 0.01% Tween at 100mg/ml in preparation for Lcn2 ELISA. Samples were vortexed for 10 minutes then centrifuged for 10 minutes at 12,000 rpm. The supernatant was collected and stored at −80°C. ELISA was performed using the Mouse Lipocalin-2/NGAL DuoSet ELISA (R&D Systems, DY1857) kit according to the manufacturer’s protocol.

### Isolation and *in vitro* stimulation of peritoneal cavity cells

Mice were euthanized and 5 ml of cold sterile PBS was injected into the peritoneal cavity via a 30G needle followed by gentle shaking to release peritoneal cavity cells. The PBS containing a single cell suspension was then retrieved via a 27G needle and immediately stored on ice. Cells were pelleted, washed, re-suspended in cold R10 medium and counted in preparation for *in vitro* stimulation. Cells were plated at 1×10^5^ cells/well in 96 well plates and stimulated with flagellins at a final concentration of 10nM for 3.5 hours at 37°C. Supernatants were harvested and Cxcl1 was measured by ELISA.

## Acknowledgments

The authors would like to acknowledge the contributions of the UAB Microbiota Culture and Analytics Resource (MCAR), the Gnotobiotic Mouse Core (GMC), and Comparative Pathology Laboratory (CPL). Research reported in this publication utilized the services of the UAB High Resolution Imaging Facility (HRIF), the UAB Mass Spectrometry/Proteomics Shared Facility (MS/PSF), and the Heflin center for Genomic Sciences, all of which receive support from the National Cancer Institute (NCI) O’Neal Comprehensive Cancer Center (OCCC) Support Grant P30 CA013148. We thank Dr. Liang Zhou (University of Florida) for providing *Tlr5*−/− colons, Dr. S Garantziotis (NIEHS) for providing *Lyz2*^Cre^.*Tlr5*^FL/FL^ mice, and L. Harrington for critical reading of the manuscript.

## Funding

National Institutes of Health grant R01 AI162736 (CLM)

National Institutes of Health grant R21 DK118386 (CLM)

National Institutes of Health grant T32 AI007051 (MSJ)

National Institutes of Health grant P01 DK071176 (COE)

Crohn’s and Colitis Foundation Career Development Award 826112 (QZ)

UAB Heersink School of Medicine Pittman Scholars Program (CLM)

## Author contributions

Conceptualization: LWD, CLM

Methodology: LWD, MSJ, CFA, KLA, GL, QZ, COE, CLM

Investigation: LWD, MSJ, JSH, CFA, ELM, KJC, BJC, QZ

Data analysis: LWD, MSJ, JSH, CFA, DH, AFR, GL, CLM

Funding acquisition: COE, CLM

Project administration: CLM

Supervision: CLM

Writing – original draft: LWD, MSJ, JSH, CFA, AFR, GL, CLM

Writing – review & editing: LWD, JSH, CFA, KLA, AFR, QZ, COE, CLM

## Data and materials availability

The RNA sequencing data for this study will be deposited upon acceptance of the manuscript. All other data needed to support the conclusions of the paper are present in the paper or the Supplementary Materials.

## Supplementary Figures

**Fig. S1:**
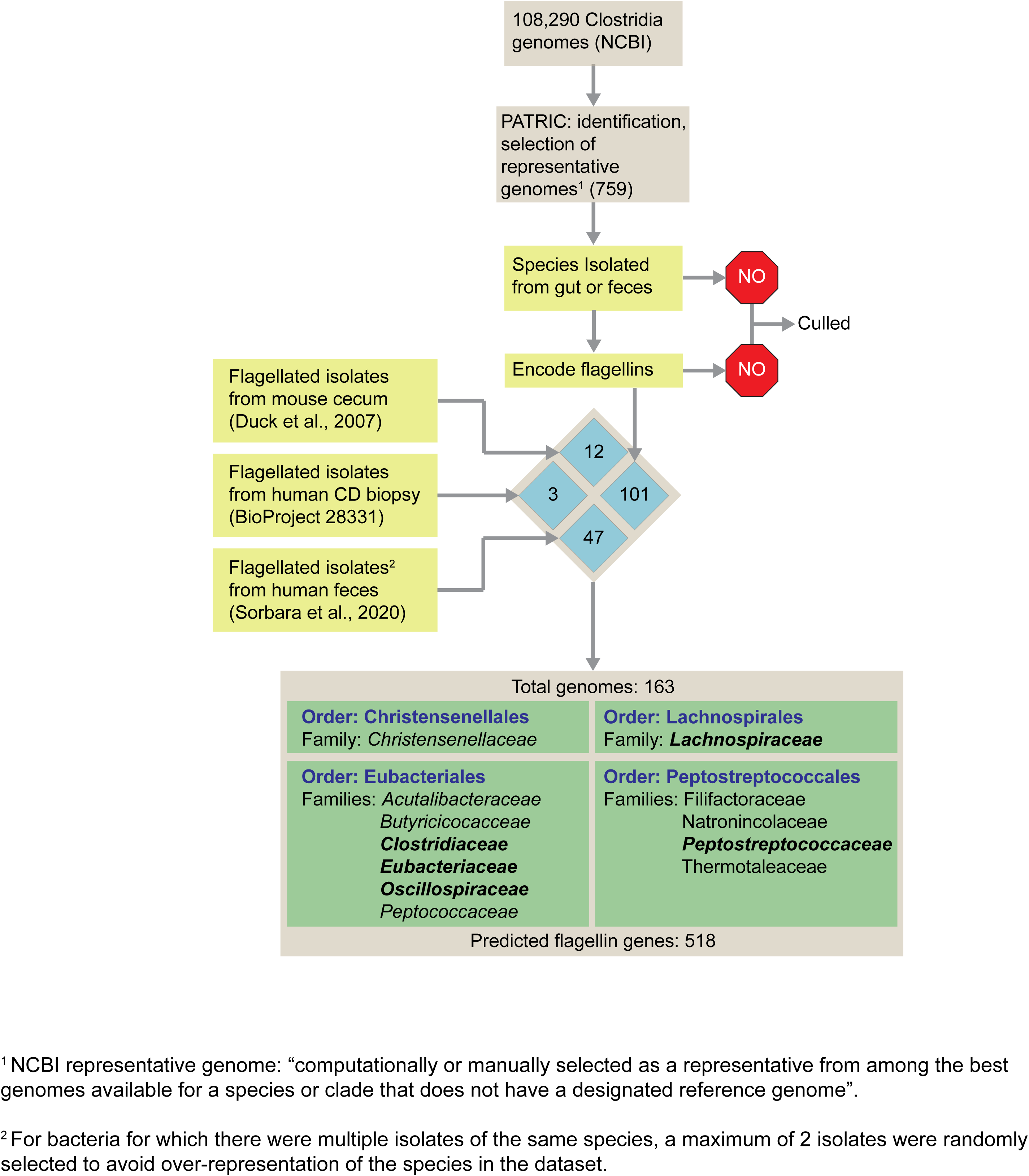
**Genome selection workflow**

**Fig. S2:**
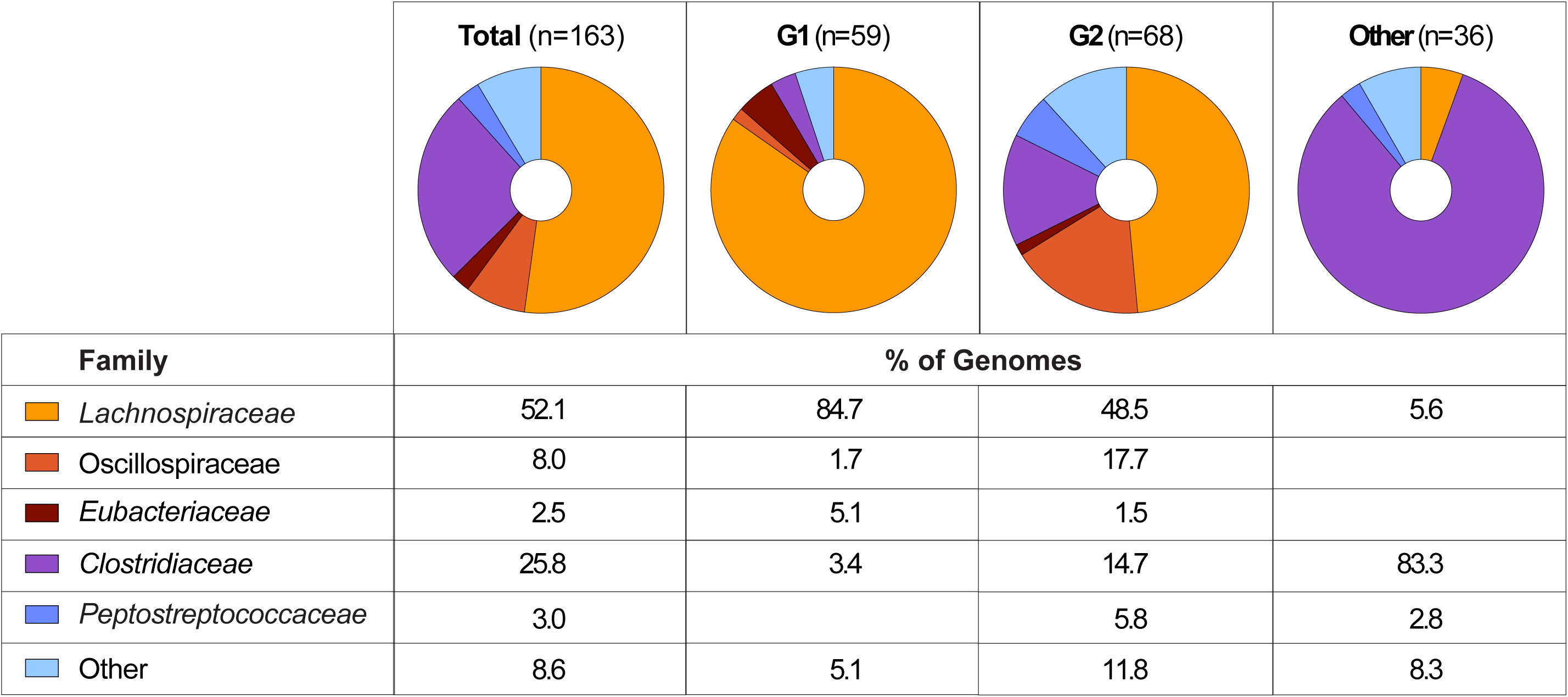
**Family distribution of G1 and G2 genomes.**

**Fig. S3.**
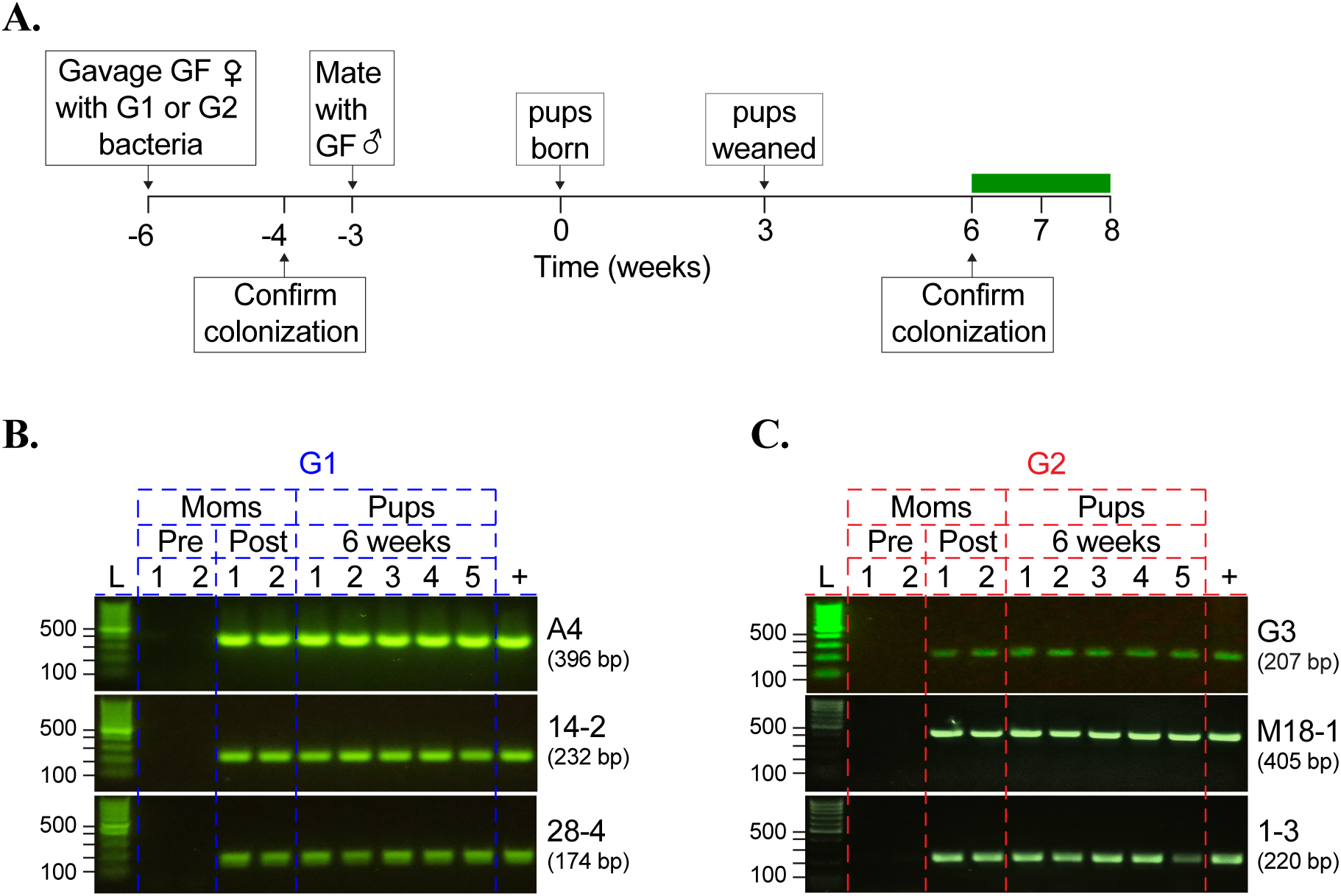
Generation and screening of gnotobiotic mice (representative workflow for a single generation of mice). (**A**) Schematic detailing the generation of gnotobiotic cohorts harboring select G1 or G2 bacteria. Adult GF mice were gavaged with G1 (*Lachnospiraceae* bacteria A4, 14-2, and 28-4) or G2 (*Anaerotruncus* sp. G3, *Lachnospiraceae* bacterium M18-1, and *Oscillobacter* sp. 1-3) bacteria at 5-6 weeks of age and mated with age-matched GF males after colonization with all bacteria was confirmed. Progeny of these matings were screened at 6 weeks of age for the presence of all organisms and used in experiments at 8 weeks of age. Representative PCR screening of feces of individual litters of (**B**) G1- and (**C**) G2-colonized progeny with maternal samples pre- and post-gavage serving as negative and positive controls, respectively.

**Fig. S4.**
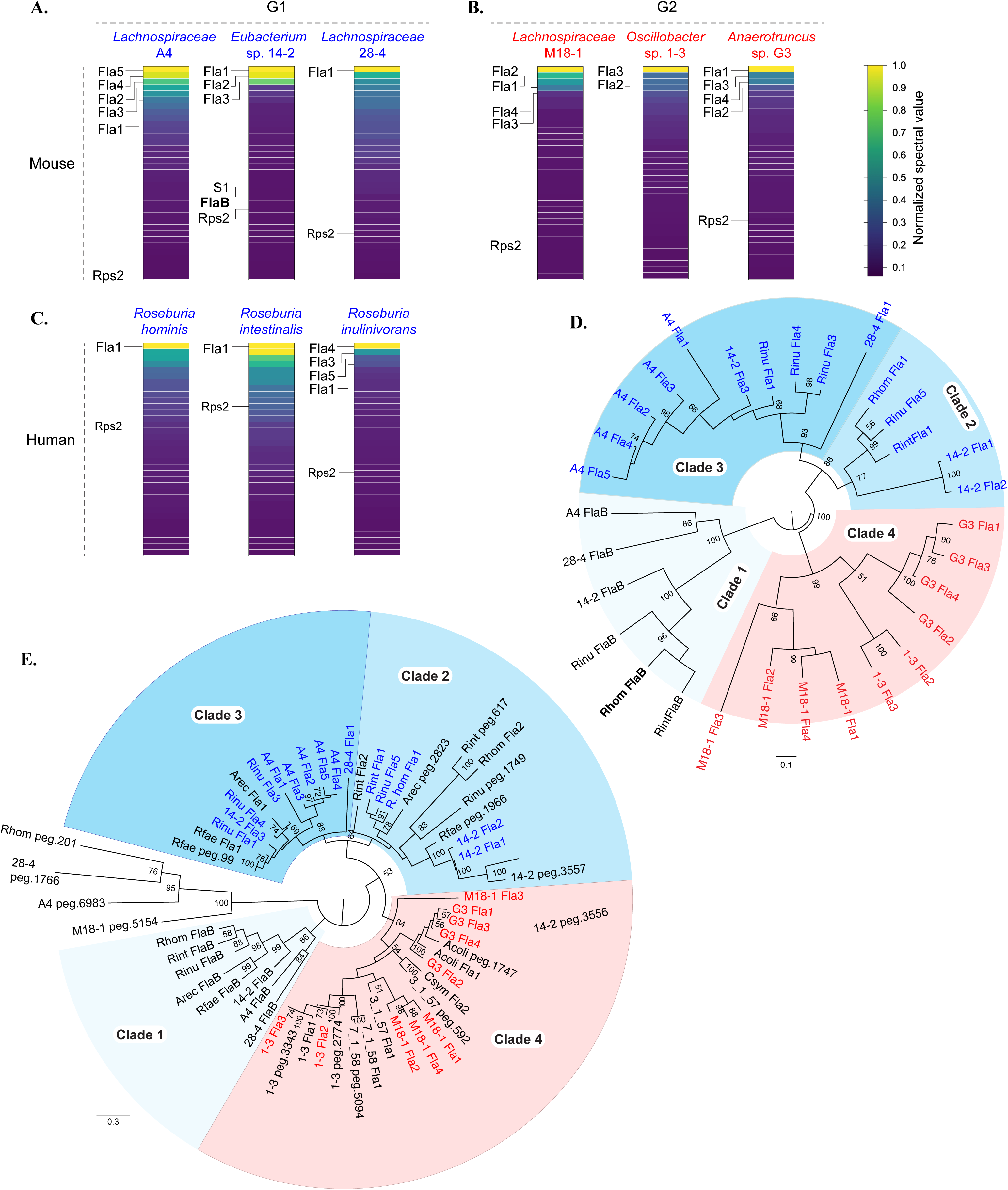
Detection and phylogenic characterization of G1- and G2-encoded flagellins. Heatmaps show normalized spectral values corresponding to the top 35 proteins identified by mass spectrometry following mechanical dissociation of (A) mouse G1 isolates used to generate gnotobiotic mice, (B) mouse G2 isolates used to generate gnotobiotic mice, and (C) select human G1 isolates of the genus Roseburia. Spectral value of each protein was normalized to that of the most abundant protein in each sample, which was set at 100%. (D) Phylogenic tree of all G1 (blue) and G2 (red) flagellins detected in A-C along with the FlaB-like flagellin (black) encoded in the respective G1 genomes. (E) Phylogenic tree of all flagellins encoded by all bacteria used in the study with detected flagellins from A-C highlighted as in D. The FlaB-like flagellin encoded in the respective G1 genomes are also indicated in Clade 1. In D-E, the Clade 1 flagellin R. hominis FlaB (Rhom FlaB) is highlighted for reference and bootstrap values of ≥ 50 are included on the trees. [Acoli: Anaerotruncus colihominis, Arec: Agathobacter rectale, Csym: Clostridium symbiosum, Rfae: Roseburia faecis, Rint: Roseburia intestinalis, Rinu: Roseburia inulinivorans, 3_1_57: Lachnospiraceae bacterium 3_1_57FAA_CT1, 7_1_58: Lachnospiraceae bacterium 7_1_58FAA].

**Fig. S5.**
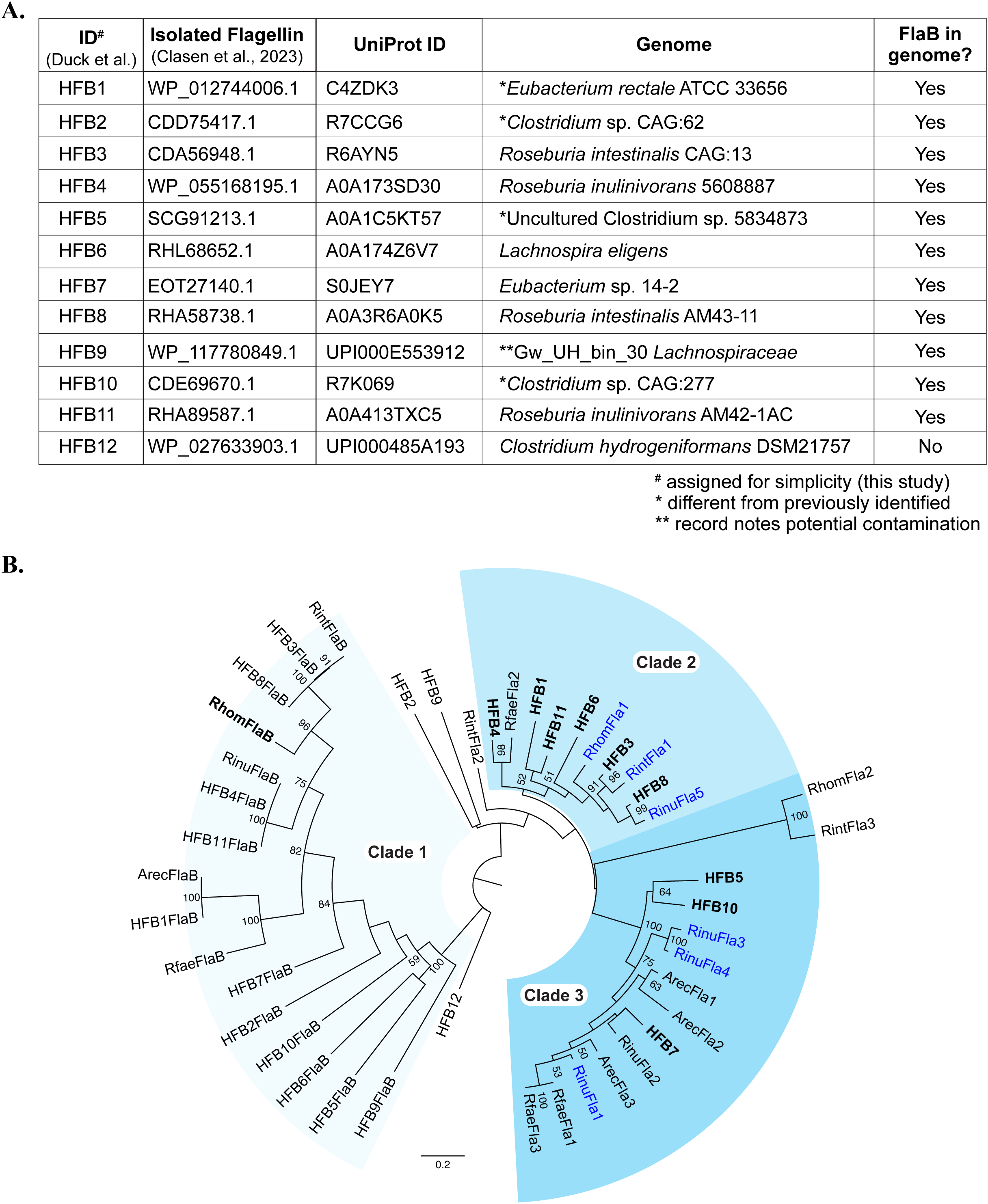
Flagellins isolated from human stool resemble G1 flagellins but do not include FlaB-like flagellins. **(A)**. Flagellins previously isolated from human stool were assigned a generic ID and protein sequences matching the published UNIPROT ID were used to identify genomes displaying a 100% match as well as any FlaB encoded \in the genome. (**B**) A phylogenetic tree of the identified flagellins along with matched FlaB flagellins encoded in the same genome. Flagellins detected as in figure 4 are included for reference. Bootstrap values ≥ 50 are shown.

**Fig. S6.**
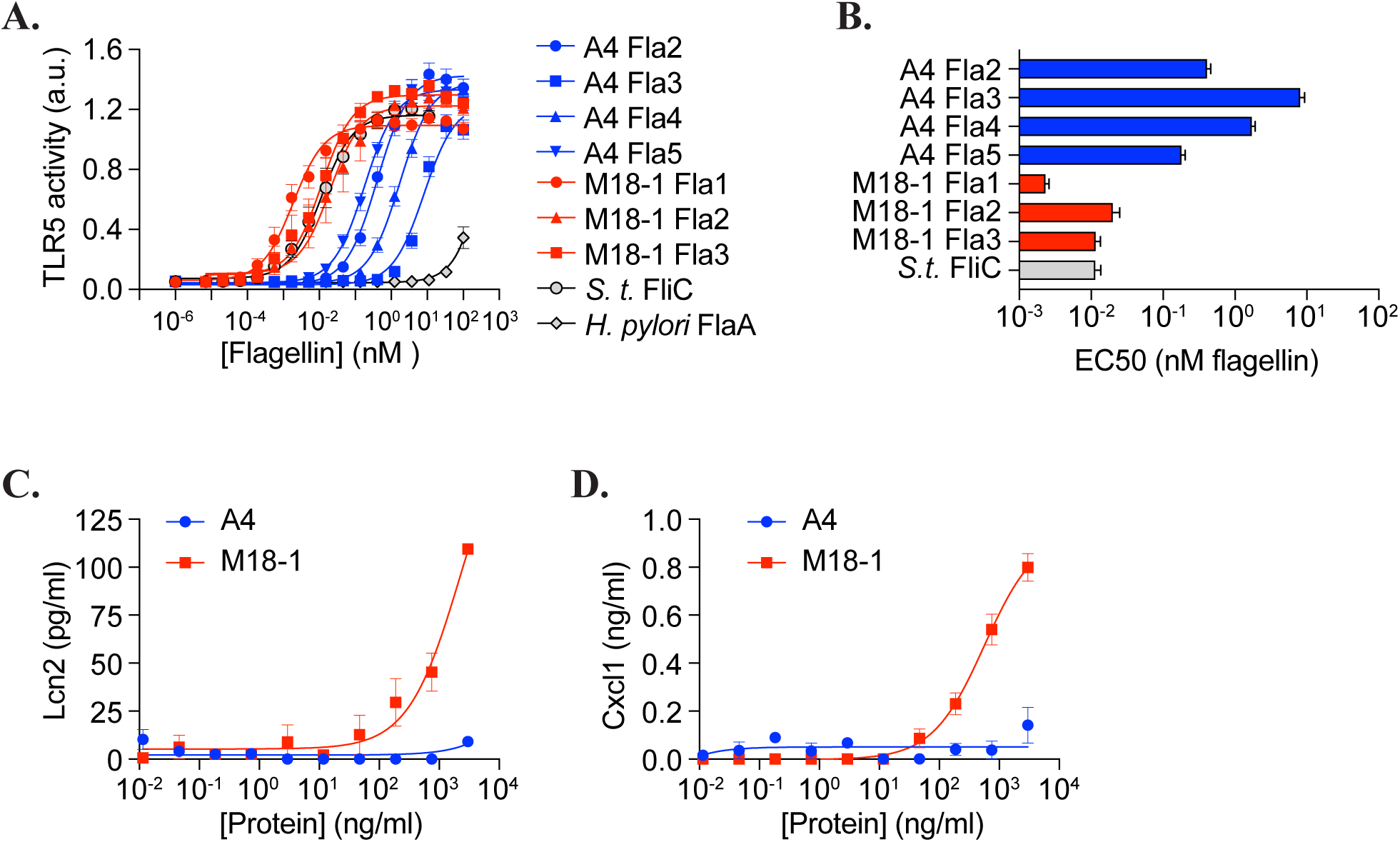
Distinct individual and collective TLR5 stimulatory capacities of Lachnospiraceae A4 and M18-1 flagellins. HEK-blue mTLR5 cells were stimulated for 16 hours with recombinant flagellins of Lachnospiraceae bacteria A4 and M18-1, representative of flagellins shown by mass spectrometry to be expressed by the respective organisms. (**A**) Individual dose response curves and (**B**) average EC50 for each individual flagellin compared to *Salmonella typhimurium* (*S.t.*) FliC and *Helicobacter pylori* (*H. pylori*) FlaA are shown. Organoid-derived epithelial monolayers were stimulated with bulk extracellular protein extracts of Lachnospiraceae bacteria A4 or M18-1 for 24 hours and (**C**) Lcn2 and (**D**) Cxcl1 in culture supernatants were measured by ELISA. All data are compiled from 2 independent experiments each with 2 replicates per condition.

**Fig. S7.**
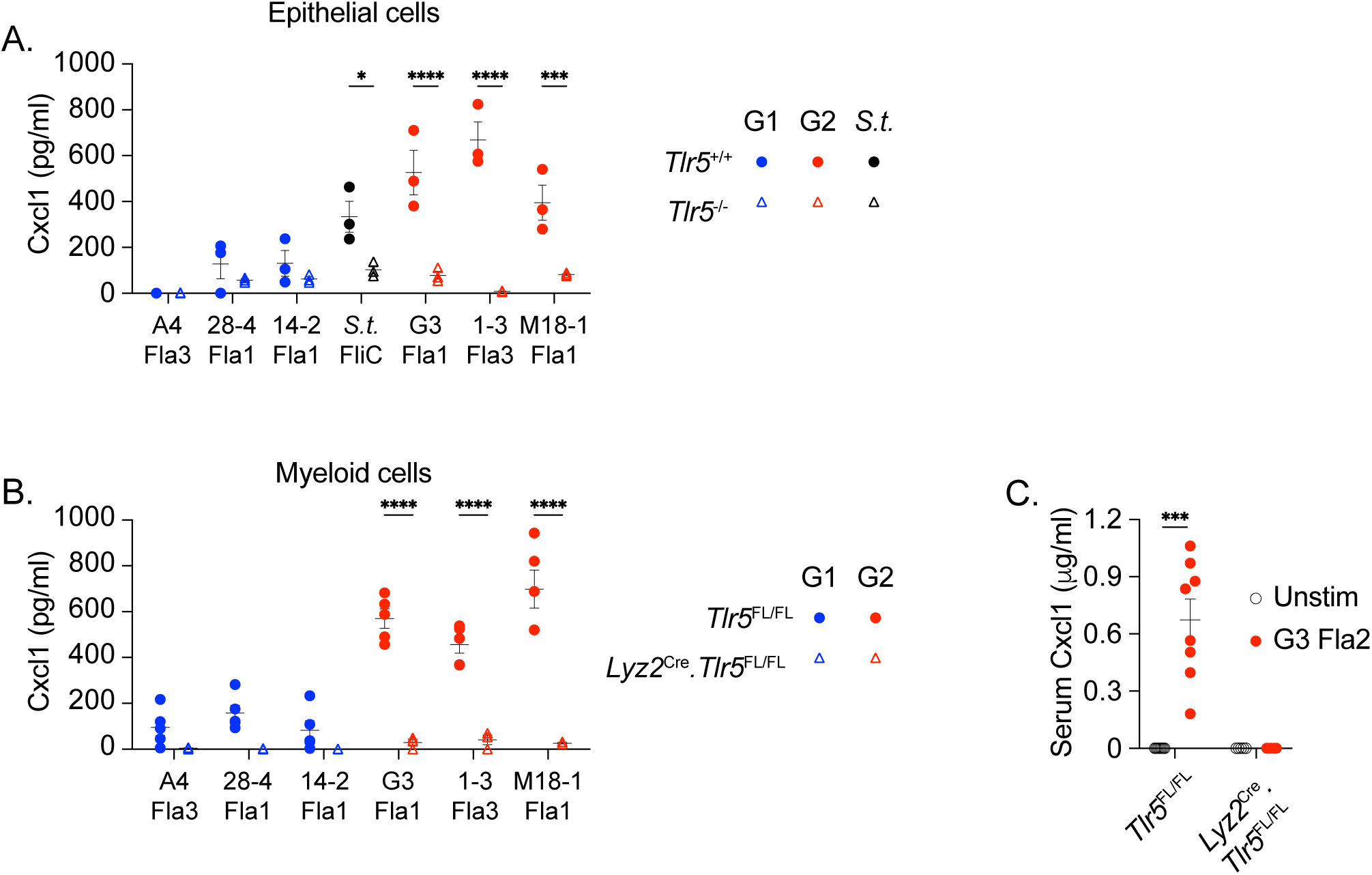
Induction of Cxcl1 by G2 flagellins is TLR5-dependent. (**A**) Colon organoid-derived epithelial monolayers prepared using WT and TLR5-deficient mice were stimulated with the indicated G1 and G2 flagellins for 24 hours. (**B**) Lymphocyte-depleted peritoneal cavity cells from *Tlr5*^FL/FL^ or *Lyz2*^Cre^.*Tlr5*^FL/FL^ mice were stimulated *in vitro* with recombinant flagellins for 3.5 hours. (**C**)*Tlr5*^FL/FL^ or *Lyz2*^Cre^.*Tlr5*^FL/FL^ mice received single intraperitoneal injections of recombinant *Anaerotruncus* G3 Fla2 and serum was collected before and 2 hours after injections. Cxcl1 in culture supernatants or serum was measured by ELISA. Data are from 1 of 2 or 3 independent experiments with similar results. Graphs show mean +/− SEM. *p<0.05, **p<0.01.

**Fig. S8.**
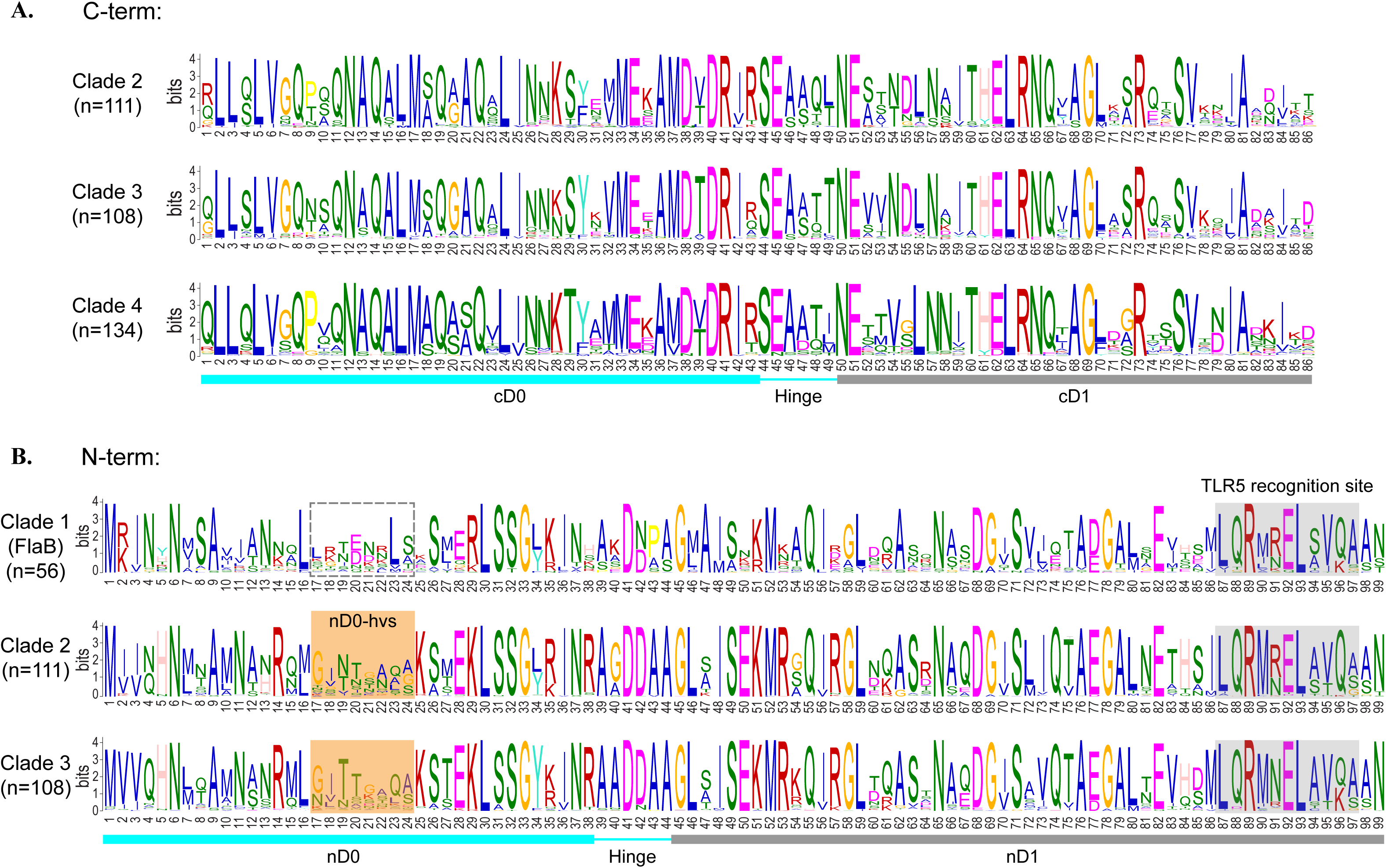
Per-residue analysis of the D0/D1 domains of flagellins by clades. Per residue information in bits were computed using MEME Suite for (**A**) the C-term end of flagellins in Clades 2, 3, and 4, and (**B**) the N-term end of flagellins in clades 1, 2, and 3. Locations of the nD0 hvs and TLR5 recognition site are indicated and numbers of flagellins used in each computation are shown in parentheses.

**Fig. S9.**
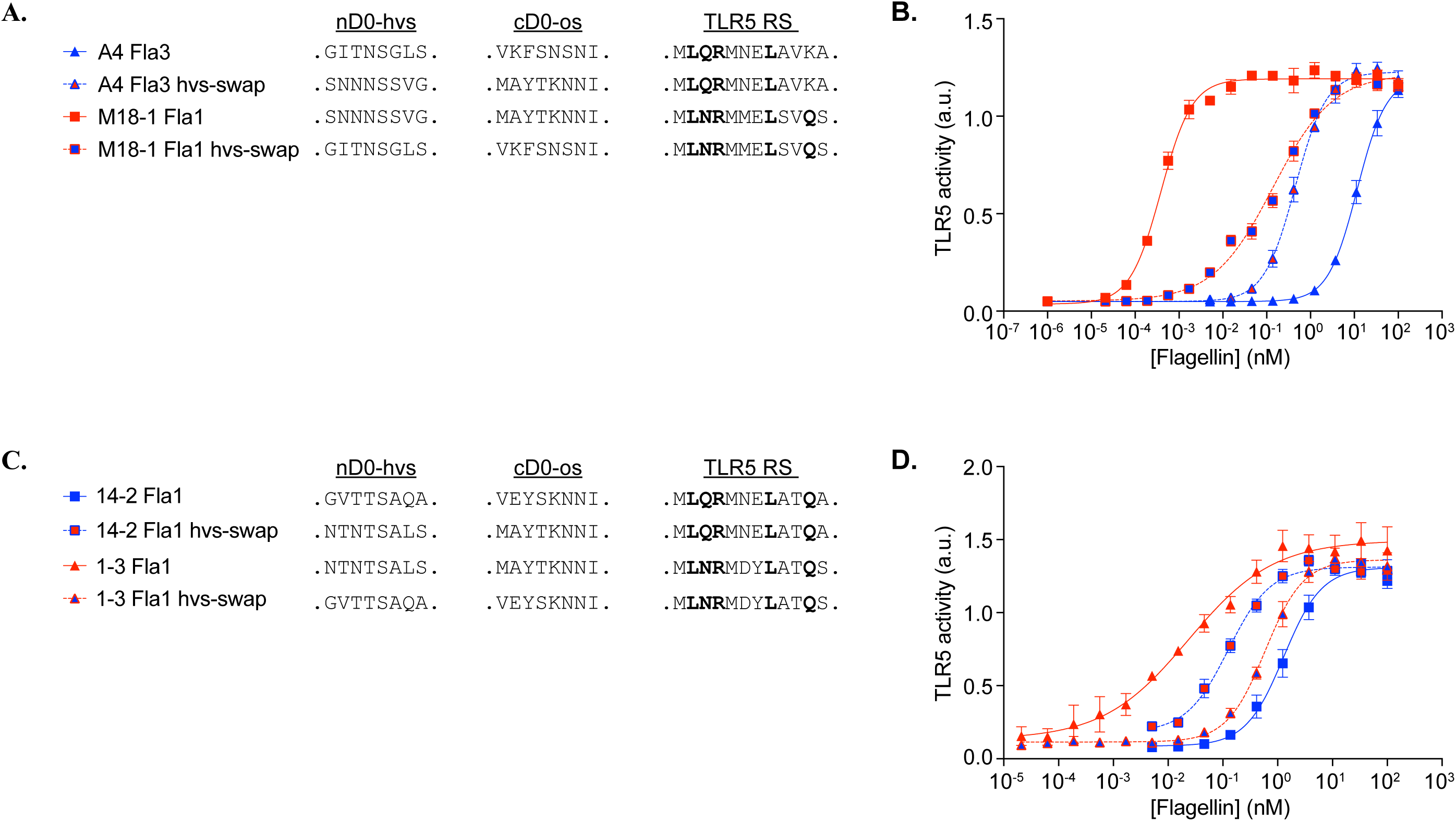
Modulation of the stimulatory capacity of G1 and G2 flagellins by altering the D0 hypervariable sequence. **(A)** Sequences of the N- and C-terminal D0 epitopes as well as the TLR5 recognition sites of the native and hvs-swapped *Lachnospiraceae* A4 Fla3 and *Lachnospiraceae* M18-1 Fla1. (**B**) Dose response curves showing TLR5 activity following 16-hour stimulation of HEK-blue mTLR5 cells with the native and hvs-swapped flagellins *Lachnospiraceae*A4 Fla3 and *Lachnospiraceae* M18-1 Fla1 flagellins. (**C**) Sequences of the N- and C-terminal D0 epitopes as well as the TLR5 recognition sites of the native and hvs-swapped *Lachnospiraceae* 14-2 Fla1 and *Oscillobacter* 1-3 Fla1. (**D**) Dose response curves showing TLR5 activity following 16-hour stimulation of HEK-blue mTLR5 cells with the native and hvs-swapped flagellins 14-2 Fla1 and 1-3 Fla1 flagellins.

## Supplementary Tables

Table S1. Genome information

Table S2. Flagellin ID and sequence

Table S3 – S11. Mass spectrometry spectral counts for select G1 and G2 bacteria examined.

Table S12. Spectral count for *Lachnospiraceae* A4 proteins detected *in vivo*.

Table S13. Sequences of primers used in the study.

## REFERENCES

1. J. Qin, R. Li, J. Raes, M. Arumugam, K. S. Burgdorf, C. Manichanh, T. Nielsen, N. Pons, F. Levenez, T. Yamada, D. R. Mende, J. Li, J. Xu, S. Li, D. Li, J. Cao, B. Wang, H. Liang, H. Zheng, Y. Xie, J. Tap, P. Lepage, M. Bertalan, J. M. Batto, T. Hansen, D. Le Paslier, A. Linneberg, H. B. Nielsen, E. Pelletier, P. Renault, T. Sicheritz-Ponten, K. Turner, H. Zhu, C. Yu, S. Li, M. Jian, Y. Zhou, Y. Li, X. Zhang, S. Li, N. Qin, H. Yang, J. Wang, S. Brunak, J. Dore, F. Guarner, K. Kristiansen, O. Pedersen, J. Parkhill, J. Weissenbach, H. I. T. C. Meta, P. Bork, S. D. Ehrlich, J. Wang, A human gut microbial gene catalogue established by metagenomic sequencing. Nature 464, 59–65 (2010).

2. N. Wadhwa, H. C. Berg, Bacterial motility: machinery and mechanisms. Nat Rev Microbiol 20, 161–173 (2022).

3. F. Hayashi, K. D. Smith, A. Ozinsky, T. R. Hawn, E. C. Yi, D. R. Goodlett, J. K. Eng, S. Akira, D. M. Underhill, A. Aderem, The innate immune response to bacterial flagellin is mediated by Toll-like receptor 5. Nature 410, 1099–1103 (2001).

4. K. L. Lightfield, J. Persson, S. W. Brubaker, C. E. Witte, J. von Moltke, E. A. Dunipace, T. Henry, Y. H. Sun, D. Cado, W. F. Dietrich, D. M. Monack, R. M. Tsolis, R. E. Vance, Critical function for Naip5 in inflammasome activation by a conserved carboxy-terminal domain of flagellin. Nat Immunol 9, 1171–1178 (2008).

5. P. V. Bulieris, N. H. Shaikh, P. L. Freddolino, F. A. Samatey, Structure of FlgK reveals the divergence of the bacterial Hook-Filament Junction of Campylobacter. Sci Rep 7, 15743 (2017).

6. B. A. Neville, P. O. Sheridan, H. M. Harris, S. Coughlan, H. J. Flint, S. H. Duncan, I. B. Jeffery, M. J. Claesson, R. P. Ross, K. P. Scott, P. W. O’Toole, Pro-inflammatory flagellin proteins of prevalent motile commensal bacteria are variably abundant in the intestinal microbiome of elderly humans. PloS one 8, e68919 (2013).

7. D. Hu, P. R. Reeves, The Remarkable Dual-Level Diversity of Prokaryotic Flagellins. mSystems 5, (2020).

8. S. J. Clasen, M. E. W. Bell, A. Borbon, D. H. Lee, Z. M. Henseler, J. de la Cuesta-Zuluaga, K. Parys, J. Zou, Y. Wang, V. Altmannova, N. D. Youngblut, J. R. Weir, A. T. Gewirtz, Y. Belkhadir, R. E. Ley, Silent recognition of flagellins from human gut commensal bacteria by Toll-like receptor 5. Sci Immunol 8, eabq7001 (2023).

9. A. M. Patterson, I. E. Mulder, A. J. Travis, A. Lan, N. Cerf-Bensussan, V. Gaboriau-Routhiau, K. Garden, E. Logan, M. I. Delday, A. G. P. Coutts, E. Monnais, V. C. Ferraria, R. Inoue, G. Grant, R. I. Aminov, Human Gut Symbiont Roseburia hominis Promotes and Regulates Innate Immunity. Front Immunol 8, 1166 (2017).

10. Z. Shen, W. Luo, B. Tan, K. Nie, M. Deng, S. Wu, M. Xiao, X. Wu, X. Meng, T. Tong, C. Zhang, K. Ma, Y. Liao, J. Xu, X. Wang, Roseburia intestinalis stimulates TLR5-dependent intestinal immunity against Crohn’s disease. EBioMedicine 85, 104285 (2022).

11. D. N. Frank, A. L. St Amand, R. A. Feldman, E. C. Boedeker, N. Harpaz, N. R. Pace, Molecular-phylogenetic characterization of microbial community imbalances in human inflammatory bowel diseases. Proc Natl Acad Sci U S A 104, 13780–13785 (2007).

12. D. Gevers, S. Kugathasan, L. A. Denson, Y. Vazquez-Baeza, W. Van Treuren, B. Ren, E. Schwager, D. Knights, S. J. Song, M. Yassour, X. C. Morgan, A. D. Kostic, C. Luo, A. Gonzalez, D. McDonald, Y. Haberman, T. Walters, S. Baker, J. Rosh, M. Stephens, M. Heyman, J. Markowitz, R. Baldassano, A. Griffiths, F. Sylvester, D. Mack, S. Kim, W. Crandall, J. Hyams, C. Huttenhower, R. Knight, R. J. Xavier, The treatment-naive microbiome in new-onset Crohn’s disease. Cell Host Microbe 15, 382–392 (2014).

13. T. Lee, T. Clavel, K. Smirnov, A. Schmidt, I. Lagkouvardos, A. Walker, M. Lucio, B. Michalke, P. Schmitt-Kopplin, R. Fedorak, D. Haller, Oral versus intravenous iron replacement therapy distinctly alters the gut microbiota and metabolome in patients with IBD. Gut 66, 863–871 (2017).

14. K. L. Alexander, Q. Zhao, M. Reif, A. F. Rosenberg, P. J. Mannon, L. W. Duck, C. O. Elson, Human Microbiota Flagellins Drive Adaptive Immune Responses in Crohn’s Disease. Gastroenterology 161, 522–535 e526 (2021).

15. G. M. Nava, H. J. Friedrichsen, T. S. Stappenbeck, Spatial organization of intestinal microbiota in the mouse ascending colon. ISME J 5, 627–638 (2011).

16. L. W. Duck, M. R. Walter, J. Novak, D. Kelly, M. Tomasi, Y. Cong, C. O. Elson, Isolation of flagellated bacteria implicated in Crohn’s disease. Inflamm Bowel Dis 13, 1191–1201 (2007).

17. M. T. Sorbara, E. R. Littmann, E. Fontana, T. U. Moody, C. E. Kohout, M. Gjonbalaj, V. Eaton, R. Seok, I. M. Leiner, E. G. Pamer, Functional and Genomic Variation between Human-Derived Isolates of Lachnospiraceae Reveals Inter- and Intra-Species Diversity. Cell Host Microbe 28, 134–146 e134 (2020).

18. .

19. C. L. Maynard, L. E. Harrington, K. M. Janowski, J. R. Oliver, C. L. Zindl, A. Y. Rudensky, C. T. Weaver, Regulatory T cells expressing interleukin 10 develop from Foxp3+ and Foxp3-precursor cells in the absence of interleukin 10. Nat Immunol 8, 931–941 (2007).

20. K. Atarashi, T. Tanoue, T. Shima, A. Imaoka, T. Kuwahara, Y. Momose, G. Cheng, S. Yamasaki, T. Saito, Y. Ohba, T. Taniguchi, K. Takeda, S. Hori, Ivanov, II, Y. Umesaki, K. Itoh, K. Honda, Induction of colonic regulatory T cells by indigenous Clostridium species. Science 331, 337–341 (2011).

21. K. Atarashi, T. Tanoue, K. Oshima, W. Suda, Y. Nagano, H. Nishikawa, S. Fukuda, T. Saito, S. Narushima, K. Hase, S. Kim, J. V. Fritz, P. Wilmes, S. Ueha, K. Matsushima, H. Ohno, B. Olle, S. Sakaguchi, T. Taniguchi, H. Morita, M. Hattori, K. Honda, Treg induction by a rationally selected mixture of Clostridia strains from the human microbiota. Nature 500, 232–236 (2013).

22. S. Basu, C. Liu, X. K. Zhou, R. Nishiguchi, T. Ha, J. Chen, M. Johncilla, R. K. Yantiss, D. C. Montrose, A. J. Dannenberg, GLUT5 is a determinant of dietary fructose-mediated exacerbation of experimental colitis. Am J Physiol Gastrointest Liver Physiol 321, G232–G242 (2021).

23. D. Jung, A. C. Fantin, U. Scheurer, M. Fried, G. A. Kullak-Ublick, Human ileal bile acid transporter gene ASBT (SLC10A2) is transactivated by the glucocorticoid receptor. Gut 53, 78–84 (2004).

24. D. Kelly, M. Kotliar, V. Woo, S. Jagannathan, J. Whitt, J. Moncivaiz, B. J. Aronow, M. C. Dubinsky, J. S. Hyams, J. F. Markowitz, R. N. Baldassano, M. C. Stephens, T. D. Walters, S. Kugathasan, Y. Haberman, N. Sundaram, M. J. Rosen, M. Helmrath, R. Karns, A. Barski, L. A. Denson, T. Alenghat, Microbiota-sensitive epigenetic signature predicts inflammation in Crohn’s disease. JCI Insight 3, (2018).

25. B. Chassaing, G. Srinivasan, M. A. Delgado, A. N. Young, A. T. Gewirtz, M. Vijay-Kumar, Fecal lipocalin 2, a sensitive and broadly dynamic non-invasive biomarker for intestinal inflammation. PloS one 7, e44328 (2012).

26. T. H. Flo, K. D. Smith, S. Sato, D. J. Rodriguez, M. A. Holmes, R. K. Strong, S. Akira, A. Aderem, Lipocalin 2 mediates an innate immune response to bacterial infection by sequestrating iron. Nature 432, 917–921 (2004).

27. B. Chassaing, R. E. Ley, A. T. Gewirtz, Intestinal epithelial cell toll-like receptor 5 regulates the intestinal microbiota to prevent low-grade inflammation and metabolic syndrome in mice. Gastroenterology 147, 1363–1377 e1317 (2014).

28. A. E. Price, K. Shamardani, K. A. Lugo, J. Deguine, A. W. Roberts, B. L. Lee, G. M. Barton, A Map of Toll-like Receptor Expression in the Intestinal Epithelium Reveals Distinct Spatial, Cell Type-Specific, and Temporal Patterns. Immunity 49, 560–575 e566 (2018).

29. S. I. Yoon, O. Kurnasov, V. Natarajan, M. Hong, A. V. Gudkov, A. L. Osterman, I. A. Wilson, Structural basis of TLR5-flagellin recognition and signaling. Science 335, 859–864 (2012).

30. V. Forstneric, K. Ivicak-Kocjan, T. Plaper, R. Jerala, M. Bencina, The role of the C-terminal D0 domain of flagellin in activation of Toll like receptor 5. PLoS Pathog 13, e1006574 (2017).

31. E. Li, C. M. Hamm, A. S. Gulati, R. B. Sartor, H. Chen, X. Wu, T. Zhang, F. J. Rohlf, W. Zhu, C. Gu, C. E. Robertson, N. R. Pace, E. C. Boedeker, N. Harpaz, J. Yuan, G. M. Weinstock, E. Sodergren, D. N. Frank, Inflammatory bowel diseases phenotype, C. difficile and NOD2 genotype are associated with shifts in human ileum associated microbial composition. PloS one 7, e26284 (2012).

32. T. Zhang, R. A. DeSimone, X. Jiao, F. J. Rohlf, W. Zhu, Q. Q. Gong, S. R. Hunt, T. Dassopoulos, R. D. Newberry, E. Sodergren, G. Weinstock, C. E. Robertson, D. N. Frank, E. Li, Host genes related to paneth cells and xenobiotic metabolism are associated with shifts in human ileum-associated microbial composition. PloS one 7, e30044 (2012).

33. J. Lloyd-Price, C. Arze, A. N. Ananthakrishnan, M. Schirmer, J. Avila-Pacheco, T. W. Poon, E. Andrews, N. J. Ajami, K. S. Bonham, C. J. Brislawn, D. Casero, H. Courtney, A. Gonzalez, T. G. Graeber, A. B. Hall, K. Lake, C. J. Landers, H. Mallick, D. R. Plichta, M. Prasad, G. Rahnavard, J. Sauk, D. Shungin, Y. Vazquez-Baeza, R. A. White, 3rd, I. Investigators, J. Braun, L. A. Denson, J. K. Jansson, R. Knight, S. Kugathasan, D. P. B. McGovern, J. F. Petrosino, T. S. Stappenbeck, H. S. Winter, C. B. Clish, E. A. Franzosa, H. Vlamakis, R. J. Xavier, C. Huttenhower, Multi-omics of the gut microbial ecosystem in inflammatory bowel diseases. Nature 569, 655–662 (2019).

34. M. Baumgart, B. Dogan, M. Rishniw, G. Weitzman, B. Bosworth, R. Yantiss, R. H. Orsi, M. Wiedmann, P. McDonough, S. G. Kim, D. Berg, Y. Schukken, E. Scherl, K. W. Simpson, Culture independent analysis of ileal mucosa reveals a selective increase in invasive Escherichia coli of novel phylogeny relative to depletion of Clostridiales in Crohn’s disease involving the ileum. ISME J 1, 403–418 (2007).

35. B. Willing, J. Halfvarson, J. Dicksved, M. Rosenquist, G. Jarnerot, L. Engstrand, C. Tysk, J. K. Jansson, Twin studies reveal specific imbalances in the mucosa-associated microbiota of patients with ileal Crohn’s disease. Inflamm Bowel Dis 15, 653–660 (2009).

36. A. Lo Presti, F. Zorzi, F. Del Chierico, A. Altomare, S. Cocca, A. Avola, F. De Biasio, A. Russo, E. Cella, S. Reddel, E. Calabrese, L. Biancone, G. Monteleone, M. Cicala, S. Angeletti, M. Ciccozzi, L. Putignani, M. P. L. Guarino, Fecal and Mucosal Microbiota Profiling in Irritable Bowel Syndrome and Inflammatory Bowel Disease. Front Microbiol 10, 1655 (2019).

37. C. Sharp, K. R. Foster, Host control and the evolution of cooperation in host microbiomes. Nat Commun 13, 3567 (2022).

38. A. M. Schoepfer, T. Schaffer, S. Mueller, B. Flogerzi, E. Vassella, B. Seibold-Schmid, F. Seibold, Phenotypic associations of Crohn’s disease with antibodies to flagellins A4-Fla2 and Fla-X, ASCA, p-ANCA, PAB, and NOD2 mutations in a Swiss Cohort. Inflamm Bowel Dis 15, 1358–1367 (2009).

39. Y. M. Ren, Z. Y. Zhuang, Y. H. Xie, P. J. Yang, T. X. Xia, Y. L. Xie, Z. H. Liu, Z. R. Kang, X. X. Leng, S. Y. Lu, L. Zhang, J. X. Chen, J. Xu, E. H. Zhao, Z. Wang, M. Wang, Y. Cui, J. Tan, Q. Liu, W. H. Jiang, H. Xiong, J. Hong, Y. X. Chen, H. Y. Chen, J. Y. Fang, BCAA-producing Clostridium symbiosum promotes colorectal tumorigenesis through the modulation of host cholesterol metabolism. Cell Host Microbe 32, 1519–1535 e1517 (2024).

40. N. N. Morgan, L. W. Duck, J. Wu, M. Rujani, P. G. Thomes, C. O. Elson, P. J. Mannon, Crohn’s Disease Patients Uniquely Contain Inflammatory Responses to Flagellin in a CD4 Effector Memory Subset. Inflamm Bowel Dis 28, 1893–1903 (2022).

41. L. Cook, D. J. Lisko, M. Q. Wong, R. V. Garcia, M. E. Himmel, E. G. Seidman, B. Bressler, M. K. Levings, T. S. Steiner, Analysis of Flagellin-Specific Adaptive Immunity Reveals Links to Dysbiosis in Patients With Inflammatory Bowel Disease. Cell Mol Gastroenterol Hepatol 9, 485–506 (2020).

42. J. Torres, F. Petralia, T. Sato, P. Wang, S. E. Telesco, R. S. Choung, R. Strauss, X. J. Li, R. M. Laird, R. L. Gutierrez, C. K. Porter, S. Plevy, F. Princen, J. A. Murray, M. S. Riddle, J. F. Colombel, Serum Biomarkers Identify Patients Who Will Develop Inflammatory Bowel Diseases Up to 5 Years Before Diagnosis. Gastroenterology 159, 96–104 (2020).

43. Q. Zhao, L. W. Duck, J. T. Killian, Jr., A. F. Rosenberg, P. J. Mannon, R. G. King, L. A. Denson, S. Kugathasan, E. N. Janoff, M. C. Jenmalm, C. O. Elson, Crohn’s Patients and Healthy Infants Share Immunodominant B Cell Response to Commensal Flagellin Peptide Epitopes. Gastroenterology 167, 1415–1428 (2024).

44. Q. Zhao, C. L. Maynard, Mucus, commensals, and the immune system. Gut Microbes 14, 2041342 (2022).

45. J. J. Gillespie, A. R. Wattam, S. A. Cammer, J. L. Gabbard, M. P. Shukla, O. Dalay, T. Driscoll, D. Hix, S. P. Mane, C. Mao, E. K. Nordberg, M. Scott, J. R. Schulman, E. E. Snyder, D. E. Sullivan, C. Wang, A. Warren, K. P. Williams, T. Xue, H. S. Yoo, C. Zhang, Y. Zhang, R. Will, R. W. Kenyon, B. W. Sobral, PATRIC: the comprehensive bacterial bioinformatics resource with a focus on human pathogenic species. Infect Immun 79, 4286–4298 (2011).

46. K. Tamura, D. Peterson, N. Peterson, G. Stecher, M. Nei, S. Kumar, MEGA5: molecular evolutionary genetics analysis using maximum likelihood, evolutionary distance, and maximum parsimony methods. Mol Biol Evol 28, 2731–2739 (2011).

47. R. C. Edgar, MUSCLE: a multiple sequence alignment method with reduced time and space complexity. BMC Bioinformatics 5, 113 (2004).

48. D. T. Jones, W. R. Taylor, J. M. Thornton, The rapid generation of mutation data matrices from protein sequences. Comput Appl Biosci 8, 275–282 (1992).

49. S. Q. Le, O. Gascuel, An improved general amino acid replacement matrix. Mol Biol Evol 25, 1307–1320 (2008).

50. N. Saitou, M. Nei, The neighbor-joining method: a new method for reconstructing phylogenetic trees. Mol Biol Evol 4, 406–425 (1987).

51. O. Gascuel, BIONJ: an improved version of the NJ algorithm based on a simple model of sequence data. Mol Biol Evol 14, 685–695 (1997).

52. FigTree Software.

53. U. Erben, C. Loddenkemper, K. Doerfel, S. Spieckermann, D. Haller, M. M. Heimesaat, M. Zeitz, B. Siegmund, A. A. Kuhl, A guide to histomorphological evaluation of intestinal inflammation in mouse models. Int J Clin Exp Pathol 7, 4557–4576 (2014).

54. D. K. Meyerholz, A. P. Beck, Principles and approaches for reproducible scoring of tissue stains in research. Lab Invest 98, 844–855 (2018).

55. H. Miyoshi, T. S. Stappenbeck, In vitro expansion and genetic modification of gastrointestinal stem cells in spheroid culture. Nat Protoc 8, 2471–2482 (2013).

56. S. H. Duncan, G. L. Hold, A. Barcenilla, C. S. Stewart, H. J. Flint, Roseburia intestinalis sp. nov., a novel saccharolytic, butyrate-producing bacterium from human faeces. Int J Syst Evol Microbiol 52, 1615–1620 (2002).

57. S. H. Duncan, R. I. Aminov, K. P. Scott, P. Louis, T. B. Stanton, H. J. Flint, Proposal of Roseburia faecis sp. nov., Roseburia hominis sp. nov. and Roseburia inulinivorans sp. nov., based on isolates from human faeces. Int J Syst Evol Microbiol 56, 2437–2441 (2006).

58. M. R. Ludwig, K. Kojima, G. J. Bowersock, D. Chen, N. C. Jhala, D. J. Buchsbaum, W. E. Grizzle, C. A. Klug, J. A. Mobley, Surveying the serologic proteome in a tissue-specific kras(G12D) knockin mouse model of pancreatic cancer. Proteomics 16, 516–531 (2016).

59. TrimGalore.

60. A. Dobin, C. A. Davis, F. Schlesinger, J. Drenkow, C. Zaleski, S. Jha, P. Batut, M. Chaisson, T. R. Gingeras, STAR: ultrafast universal RNA-seq aligner. Bioinformatics 29, 15–21 (2013).

61. S. Anders, P. T. Pyl, W. Huber, HTSeq--a Python framework to work with high-throughput sequencing data. Bioinformatics 31, 166–169 (2015).

62. M. D. Robinson, D. J. McCarthy, G. K. Smyth, edgeR: a Bioconductor package for differential expression analysis of digital gene expression data. Bioinformatics 26, 139–140 (2010).

